# Gardos potassium channel amplifies PIEZO1-TMEM16F coupling in red blood cells

**DOI:** 10.64898/2026.05.28.728404

**Authors:** Yui Chun Serena Wan, Martha Delahunty, Grace Lee, Sanjay Khandelwal, Gowthami M. Arepally, Marilyn J. Telen, Huanghe Yang

## Abstract

In red blood cells (RBCs), the mechanosensitive channel PIEZO1 provides an upstream Ca^2+^ signal that activates TMEM16F, a Ca^2+^-activated phospholipid scramblase (CaPLSase) responsible for phosphatidylserine (PS) externalization. However, limited PIEZO1-mediated Ca²⁺ entry and the relatively low Ca^2+^ sensitivity of TMEM16F suggest the need for signal amplification. Here, we identify the Gardos (KCNN4) Ca^2+^-activated K⁺ channel as a critical amplifier of the PIEZO1-TMEM16F axis. Gardos activation induces membrane hyperpolarization, thereby increasing the driving force for Ca^2+^ entry and enhancing TMEM16F activation and phospholipid scrambling. Gardos-mediated amplification also contributes to excessive PS externalization in sickle cell disease (SCD) and hereditary xerocytosis (HX) RBCs, and functional disruption of Gardos-mediated K^+^ efflux attenuates this response. These findings demonstrate Gardos as a critical amplifier of RBC mechanotransduction and highlight Gardos as a potential therapeutic target for mitigating pathogenic PS exposure.

**HIGHLIGHTS:**

- Gardos amplifies PIEZO1-mediated Ca²⁺ influx to promote TMEM16F-dependent phosphatidylserine exposure in healthy and diseased RBCs.
- Unexpected effects of some Gardos inhibitors may complicate the use in hematologic diseases.

## INTRODUCTION

Phosphatidylserine (PS) exposure on the outer leaflet of the plasma membrane is a tightly regulated process that serves as a key signal for coagulation, phagocytic clearance, and cell-cell interactions^1–8^. In blood cells, PS exposure is strongly associated with pathological conditions, including thrombosis, hemolysis, and vascular dysfunction, and is a hallmark and pathological driver of blood disorders such as sickle cell disease (SCD) and hereditary xerocytosis (HX)^9–12^. Excessive PS exposure on RBCs promotes procoagulant activity, microparticle shedding, and endothelial adhesion, thereby contributing to vaso-occlusion and cardiovascular complications in these diseases^11–21^.

At the molecular level, recent advances in phospholipid transport have identified TMEM16F as the long-sought Ca²⁺-activated phospholipid scramblase (CaPLSase) responsible for Ca^2+^-dependent PS exposure in platelets and RBCs^13,22,23^. TMEM16F-mediated PS externalization is essential for platelet procoagulant activity, and loss-of-function mutations cause Scott syndrome, a rare hereditary bleeding disorder^13,22–24^. Conversely, increased TMEM16F activity is observed in SCD and HX RBCs, leading to pathological PS exposure and PS-mediated complications such as thrombosis, anemia, and vaso-occlusion^10,11^. We recently identified a mechanochemical coupling mechanism in which the mechanosensitive, Ca^2+^-permeable ion channel PIEZO1 provides the upstream Ca^2+^ signal that activates TMEM16F in RBCs (Fig. 1A). This PIEZO1-TMEM16F axis constitutes a key pathway underlying excessive PS exposure in SCD and HX^10,11^. Consistent with this model, pharmacological inhibition of PIEZO1 attenuates TMEM16F activation and PS externalization^10,11,25^. These findings suggest that targeting the PIEZO1-TMEM16F axis represents a promising therapeutic strategy for mitigating RBC-driven thrombosis and vascular complications.

**Figure 1.**
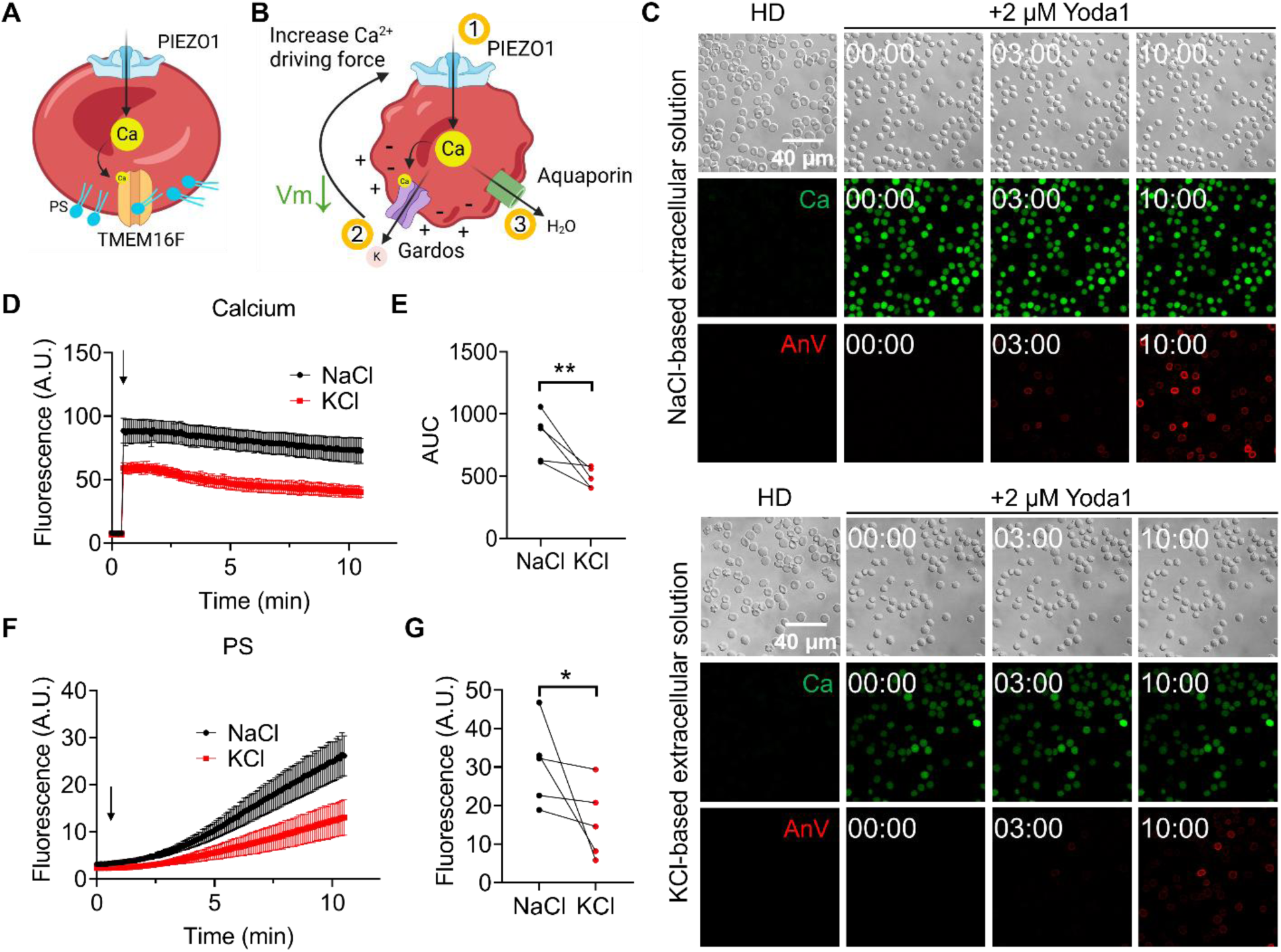
Collapsing the K^+^ gradient reduces Ca^2+^ influx and PS exposure in RBCs. (A) Illustration of the current model of PIEZO1-TMEM16F coupling in mediating PS exposure in RBCs. (B) Illustration of the current model of Gardos-mediated RBC changes downstream of PIEZO1 activation. PIEZO1-mediated Ca^2^ ^+^ influx activates Gardos, leading to K^+^ efflux, which drives membrane hyperpolarization and, in turn, increases the Ca^2+^ driving force through PIEZO1. (C) Representative images of simultaneous Ca^2+^ and PS time-lapse confocal imaging of healthy donor (HD) RBCs in NaCl- (top) and KCl-based (bottom) extracellular solutions. RBCs were stimulated with 2 µM Yoda1, and the morphology, Ca^2+^ levels, and PS exposure were monitored for 10 minutes after the stimulation. Ca^2+^ levels were measured by Calbryte fluorescence dye, and PS exposure was monitored using fluorophore-conjugated Annexin V (AnV). (D) Summary of Ca^2+^ imaging results, measured by Calbryte dye, of HD RBCs from five participants in NaCl- and KCl-based solutions. The arrow indicates stimulation of 2 µM Yoda1. The error bars indicate ± SEM. (E) The area under the curve (AUC) analysis of Yoda1-induced Ca^2+^ levels of HD RBCs in NaCl- and KCl-based solutions from panel D. The connected dots indicate the same HD participant; ** two-sided t-test, *P* = 0.0076 (n = 5). (F) Summary of PS exposure results of HD RBCs from five participants in NaCl- and KCl-based solutions. The arrow indicates stimulation of 2 µM Yoda1. (G) Statistical analysis of the endpoint AnV fluorescence of HD RBCs in NaCl- and KCl-based solutions from panel F. The connected dots indicate the same HD participant; * two-sided t-test, *P* = 0.0498 (n = 5).

Despite this progress, direct therapeutic targeting of this pathway remains challenging. Although TMEM16F deficiency abolishes Ca^2+^-induced PS exposure^11,22,23^, the development of selective TMEM16F inhibitors has proven difficult^26–28^. Alternatively, partial inhibition of PIEZO1 using agents such as GsMTx4, benzbromarone, or cannabidiol reduces PS exposure in SCD and HX RBCs^10,11,25^. However, none of these compounds are clinically approved or sufficiently selective, underscoring the need to further dissect the regulatory mechanisms governing PIEZO1-TMEM16F coupling and to identify alternative therapeutic targets.

A key unresolved question has been how relatively limited Ca^2+^ influx through PIEZO1 is sufficient to activate TMEM16F. PIEZO1 is sparsely expressed in RBCs (∼100 channels per cell) and exhibits rapid inactivation kinetics upon activation^29–31^, constraining the magnitude and duration of Ca^2+^ entry. In contrast, TMEM16F has relatively low Ca^2+^ sensitivity, requiring sustained micromolar Ca^2+^ elevations for activation^22,32–36^. These properties suggest that an amplification mechanism is required to efficiently couple PIEZO1 activation to TMEM16F-mediated lipid scrambling.

The Gardos channel (encoded by *KCNN4*; also known as KCa3.1, SK4, or IK) is a Ca^2+^-activated K⁺ channel and a well-established downstream effector of PIEZO1-mediated Ca^2+^ entry in RBCs^37,38^ (Fig. 1B). Notably, Gardos channels are activated by submicromolar-to-low-micromolar Ca^2+^, substantially lower than those typically required for TMEM16F activation^39–43^. Activation of Gardos channels leads to K⁺ efflux and membrane hyperpolarization, thereby increasing the electrochemical driving force for Ca^2+^ entry^38,44–47^. Based on these properties, we hypothesized that the Gardos channel functions as a critical amplifier that establishes the efficiency of PIEZO1-TMEM16F coupling. In this study, we tested this hypothesis using heterologous expression systems, erythroleukemia cell lines, and primary RBCs and established a PIEZO1-Gardos-TMEM16F signaling axis. We demonstrate that PIEZO1 activation leads to Gardos-mediated membrane hyperpolarization, amplifying Ca^2+^ influx and promoting TMEM16F activation and PS exposure. In addition, we uncover unexpected off-target effects of commonly used Gardos inhibitors, highlighting the need for more selective pharmacological tools. Together, our findings define a previously unrecognized amplification mechanism in RBC mechanotransduction and suggest that targeting the Gardos channel may provide a novel therapeutic strategy to attenuate pathogenic PS exposure in blood disorders such as SCD and HX.

## METHODS

### Human blood samples

All human studies were approved by the institutional review board at Duke University (IRB# Pro00007816, Pro00109511, and Pro00104933). Written informed consents were obtained from all participants. The RBCs were isolated as previously described with minor modifications^11^. Briefly, whole blood was collected in acid-citrate-dextrose (ACD) buffer. To avoid mechanical stimulation, 500 µL of whole blood in ACD was mixed with 1 mL of DPBS and centrifuged at 100 x g for 6 minutes. After aspirating the supernatant, the RBC pellets were used immediately for experiments. Whole blood samples were stored at 4 °C and RBCs were freshly isolated before each experiment.

### Flow cytometry measurement of PS exposure

Flow cytometry measurement of RBC PS exposure was performed as described previously^11^ with the modification that packed RBCs with a final 1:2000 dilution were added into 200 µL Hank’s Balanced Salt Solution (HBSS), NaCl-based solution, or KCl-based solution containing a 1:125 dilution of Annexin V (AnV) as indicated for the experiment. The PIEZO1 agonist Yoda1 was then added to the solution to reach a final concentration of 1 or 2 µM and incubated at room temperature for 10 minutes.

For K562 cells, the cells were first collected and pelleted with centrifugation at 500 x g for 5 min. The cells were then re-diluted to 5 × 10^6^ cells/mL in HBSS and kept on ice. 20 µL of cell dilution was resuspended in 180 µL of AnV solution (1:125 in HBSS). Cells were stimulated with 0.5 µM Yoda1 for 10 minutes at room temperature. Analysis was performed using the FlowJo Software (Waters Biosciences, Milford, MA).

### Fluorescence microscopy

For real-time Ca^2+^ measurements of cell lines, the cells were loaded with Fura2 fluorescent Ca^2+^ dye for 45 minutes in HBSS. The cells were then stimulated with Yoda1 with the desired concentrations diluted in HBSS. Using an Olympus IX83 fluorescence microscope, the cells were excited at 340 and 380 nm (pE-340^FURA^, CoolLED), and the emission of the cells was measured in real-time at 510 nm. Ca^2+^ levels were quantified as the 340/380 excitation ratio, with emission collected at 510 nm. For PS measurements, the imaging solution contained 1:125 dilution of fluorophore-conjugated Annexin V (AnV) in HBSS. The cells were then stimulated with 1 µM Yoda1 in HBSS also containing AnV. The PS exposure was then monitored for 10 minutes at room temperature in real-time with the wavelength according to the fluorophore used. For experiments using Senicapoc or clotrimazole, the inhibitors were included in both the initial imaging solution and the stimulation solution. Images were analyzed with ImageJ (version 1.54f). For HEK293T cell experiments involving overexpression, only cells that expressed the proteins of interest were selected for data analysis.

Fluorescence microscopy of RBCs was performed as previously described^11^. Briefly, RBCs were seeded on uncoated coverslips in NaCl-based extracellular solution (140 mM NaCl, 5 mM KCl, 2 mM MgCl_2_, 10 mM HEPES, and 2 mM CaCl_2_ at pH 7.4) containing 1 µM Calbryte 520 for 10 minutes at 37°C. The RBCs were then imaged in 1:125 of fluorophore-conjugated AnV in NaCl-based or KCl-based extracellular solution (147 mM KCl, 2 mM MgCl_2_, 10 mM HEPES, and 2 mM CaCl_2_ at pH 7.4). Yoda1 diluted in the respective NaCl- or KCl-based solution was added at a 1:2 volume ratio to achieve a final concentration of 2 µM. The Ca^2+^ and the PS exposure levels were simultaneously monitored for 10 minutes at room temperature in real-time with wavelengths according to the fluorophore used.

### Statistical analysis

All statistical comparisons were performed using GraphPad Prism (GraphPad, Boston, MA). Unpaired two-tailed Student’s t-tests were used when comparing two groups. One-way ANOVA tests were used for multiple comparisons. Sample sizes (n) and *P* values are indicated in the corresponding figure legends.

## RESULTS

### K^+^ gradient regulates PIEZO1-TMEM16F coupling in RBCs

Under physiological conditions, a high intracellular and low extracellular K^+^ concentration establishes a steep electrochemical gradient that drives K^+^ efflux. Activation of the Gardos channel promotes K^+^ efflux and membrane hyperpolarization in RBCs, thereby increasing the driving force for Ca^2+^ entry through Ca^2+^-permeable channels^38^ (Fig. 1B). We therefore hypothesized that replacing extracellular Na^+^ with K^+^ (i.e., switching from a NaCl- to a KCl-based solution) would collapse the K^+^ gradient, abolish Gardos-mediated K^+^ efflux and membrane hyperpolarization, and consequently attenuate Ca^2+^ influx through PIEZO1, thereby limiting TMEM16F activation and PS exposure.

To test this, we stimulated healthy donor (HD) RBCs with 2 µM Yoda1, a PIEZO1 agonist^48^, and simultaneously monitored Ca^2+^ and PS using confocal microscopy. Consistent with our previous studies^10,11^, Yoda1 elicited a rapid increase in intracellular Ca^2+^ followed by robust PS exposure in the NaCl-based solution (Fig. 1C-G). In contrast, in the KCl-based solution, Yoda1 still triggered rapid Ca^2+^ influx, but the overall magnitude of Ca^2+^ entry was significantly reduced (Fig. 1C-E), indicating that the loss of the K^+^ gradient reduces the driving force for Ca^2+^ entry through PIEZO1. Correspondingly, PS exposure was significantly reduced when the K^+^ gradient was diminished (Fig. 1C, F, G), suggesting that a K^+^ conductance downstream of PIEZO1 activation plays a critical role in facilitating Ca^2+^-activated lipid scrambling.

We next examined whether the K^+^ gradient similarly regulates Ca^2+^ entry and PS exposure in diseased RBCs. Similar to HD RBCs, Yoda1-induced Ca^2+^ influx and PS externalization in SCD (Fig. 2A-F) and HX RBCs (Fig. 2G-L) were significantly reduced in the KCl-based extracellular solution compared with the NaCl-based solution. Together, results from the HD, SCD, and HX RBCs demonstrated that a K^+^ conductance downstream of PIEZO1 is required to amplify PIEZO1-mediated Ca^2+^ influx and subsequent TMEM16F activation in healthy and diseased RBCs.

**Figure 2.**
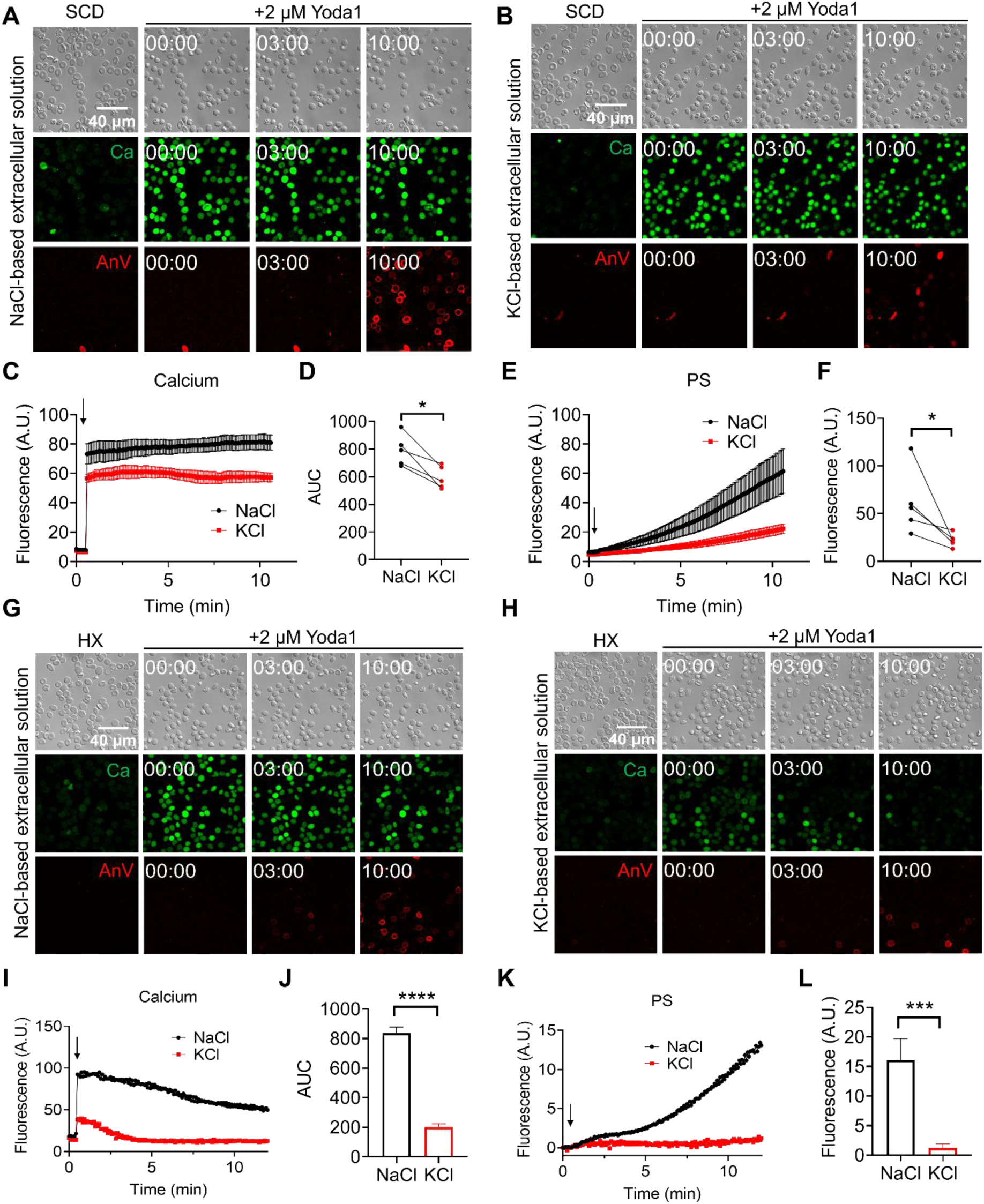
Abolishing the K^+^ gradient reduces Ca^2+^ influx and PS exposure in disease RBCs. (A-B) Representative images of simultaneous Ca^2+^ and PS imaging of SCD RBCs in NaCl- (A) and KCl-based (B) extracellular solutions before and after 10 minutes of 2 µM Yoda1 stimulation. (C) Summary of Ca^2+^ imaging results, measured by Calbryte dye, of SCD RBCs from five participants in NaCl- and KCl-based solutions. The arrow indicates stimulation of 2 µM Yoda1. The error bars indicate ± SEM. (D) The AUC analysis of Yoda1-induced Ca^2+^ levels of SCD RBCs in NaCl- and KCl-based solutions. The connected dots indicate the same SCD participant; * two-sided t-test, *P* = 0.0139 (n = 5). (E) Summary of PS exposure results of SCD RBCs from five participants in NaCl- and KCl-based solutions. The arrow indicates stimulation of 2 µM Yoda1. (F) Statistical analysis of the endpoint AnV fluorescence of SCD RBCs in NaCl- and KCl-based solutions. The connected dots indicate the same SCD participant; * two-sided t-test, *P* = 0.0359 (n = 5). (G-H) Representative images of simultaneous Ca^2+^ and PS imaging of HX RBCs from one participant in NaCl-based (G) and KCl-based (H) extracellular solutions before and after 10 minutes of 2 µM Yoda1 stimulation. (I) Summary of Ca^2+^ imaging results of HX RBCs from one participant in NaCl- and KCl-based solutions. The arrow indicates stimulation of 2 µM Yoda1. (J) The AUC analysis of Yoda1-induced Ca^2+^ levels of HX RBCs in NaCl- and KCl-based solutions. **** two-sided t-test, *P* < 0.0001 (n = 50 cells for NaCl condition; n = 40 cells for KCl condition). (K) Summary of PS exposure results of HX RBCs from one participant in NaCl- and KCl-based solutions. The arrow indicates stimulation of 2 µM Yoda1. (L) Statistical analysis of the endpoint AnV fluorescence of HX RBCs in NaCl- and KCl-based solutions. *** two-sided t-test, *P* = 0.0006, n = 50 cells for NaCl condition; n = 40 cells for KCl condition).

### Gardos amplifies PIEZO1-TMEM16F coupling in HEK293T cells

Given that Gardos is responsible for the Ca^2+^-activated K⁺ conductance in RBCs^37,49–51^, we next tested whether it amplifies PIEZO1-TMEM16F coupling by reconstituting these components in HEK293T cells. We first used whole-cell current-clamp recordings to monitor membrane potential changes in response to Yoda1 (Fig. 3A, left). HEK293T cells stably expressing PIEZO1 exhibited a small but detectable membrane depolarization upon Yoda1 stimulation (Fig. 3Ai, B), consistent with the nonselective cation permeability of PIEZO1^29^. In contrast, co-expression of Gardos with PIEZO1 resulted in robust membrane hyperpolarization toward the K⁺ equilibrium potential (∼-80 mV) following Yoda1 application (Fig. 3Aii, B). These results demonstrate that PIEZO1-mediated Ca^2+^ entry rapidly activates Gardos, whose opening drives membrane hyperpolarization.

**Figure 3.**
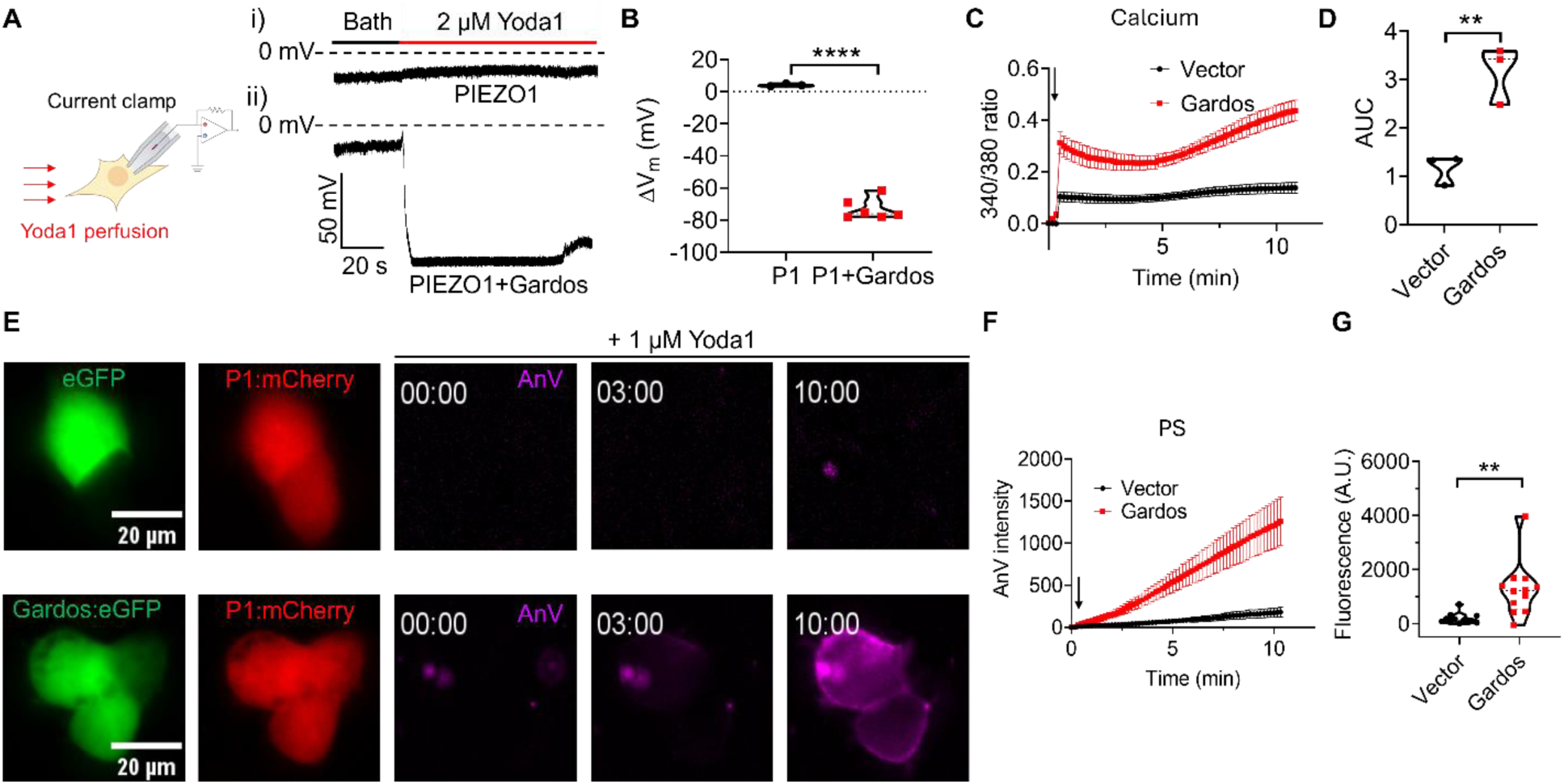
Gardos amplifies PIEZO1-TMEM16F coupling by enhancing Ca^2+^ influx through heterologously expressed PIEZO1 in HEK293T cells. (A) Illustration of HEK293T cell whole-cell patch clamping configuration with Yoda1 perfusion. Representative current clamp measurements of membrane potentials of HEK293T cells overexpressing PIEZO1 (i) or PIEZO1+Gardos (ii). The cells were first perfused with bath solution (black), followed by Yoda1 solution (red). (B) Statistical analysis of the differences between the membrane potential with and without Yoda1 perfusion. The membrane potential differences (ΔV_m_) were calculated by subtracting the membrane potential after Yoda1 perfusion by the membrane potential under bath solution perfusion. The membrane potential of HEK293T cells overexpressing PIEZO1+Gardos (n = 6) was compared to the PIEZO1-only condition (n = 3); **** two-sided t-test *P* < 0.0001. (C) Fura2 Ca^2+^ fluorescence imaging of HEK293T cells stably expressing PIEZO1 transfected with vector control or Gardos. The arrow indicates stimulation of 1 µM Yoda1, and the Ca^2+^ influx levels were monitored for 10 minutes after stimulation. The error bars indicate ± standard error of the mean (SEM) of 3 replicates of each sample. (D) The AUC analysis of Ca^2+^ levels from panel C. A violin plot represented the AUC results (n = 3); ** two-sided t-test *P* = 0.0067. (E) Representative images of HEK293T cells exposing PS as detected by AnV (magenta). The top panel represents PIEZO1 stable cells (red) overexpressing eGFP vector (green), and the bottom panel represents PIEZO1 stable cells (red) overexpressing Gardos (green). (F) Summary of real-time PS exposure of HEK293T PIEZO1 stable cells overexpressing vector control or Gardos. The arrow indicates stimulation of 1 µM Yoda1. The error bars indicate ± SEM of 12 replicates of each sample. (G) Statistical analysis of endpoint fluorescence values of panel F; ** two-sided t-test, *P* = 0.0014 (n = 12).

Fura-2 Ca^2+^ imaging further revealed that co-expression of PIEZO1 and Gardos significantly enhanced and prolonged intracellular Ca^2+^ elevation compared with PIEZO1 expression alone (Fig. 3C-D). Because membrane hyperpolarization increases the electrochemical driving force for Ca^2+^ entry, these findings support a positive-feedback mechanism in which Gardos activation downstream of PIEZO1 amplifies Ca^2+^ influx through PIEZO1.

We next asked whether this enhanced Ca^2+^ influx leads to increased PS exposure via endogenous TMEM16F in HEK293T cells. Consistent with this prediction, cells co-expressing PIEZO1 and Gardos exhibited significantly greater PS exposure than cells expressing PIEZO1 alone following Yoda1 stimulation (Fig. 3E-G). Together, these results establish that Gardos amplifies PIEZO1-TMEM16F coupling by enhancing PIEZO1-mediated Ca^2+^ entry and subsequent TMEM16F-dependent phospholipid scrambling.

### Gardos channelopathy S314P promotes PIEZO1-TMEM16F coupling in HEK293T cells

Gain-of-function (GOF) *KCNN4*/Gardos channelopathy causes dehydrated hereditary stomatocytosis or Gardos-mediated HX, an autosomal dominant hemolytic anemia characterized by excessive K^+^ loss and RBC dehydration^39,52–57^. We hypothesized that such GOF mutations would potentiate Ca^2+^ influx through PIEZO1, thereby enhancing TMEM16F activation and subsequent PS exposure. To test this, we overexpressed the Gardos channelopathy variant S314P, which increases the Ca^2+^ sensitivity of Gardos^39^, in HEK293T cells stably expressing PIEZO1. We then monitored Ca^2+^ influx and PS exposure following stimulation with a low concentration of Yoda1 (0.2 µM). The weak Yoda1 stimulation induced modest Ca^2+^ entry and PS exposure in cells co-expressing PIEZO1 and wild-type (WT) Gardos (Fig. 4A, C–F). However, both Ca^2+^ influx and PS externalization were significantly enhanced in cells expressing the S314P mutant compared with WT Gardos (Fig. 4B–F). Together, these results demonstrate that increased Gardos activity, either through overexpression of WT (Fig. 3) or disease-associated GOF mutation (Fig. 4), potentiates PIEZO1-TMEM16F coupling. These findings provide mechanistic insight into the pathophysiology of Gardos channelopathy by linking enhanced Gardos activity to amplified Ca^2+^ signaling and excessive PS exposure.

**Figure 4.**
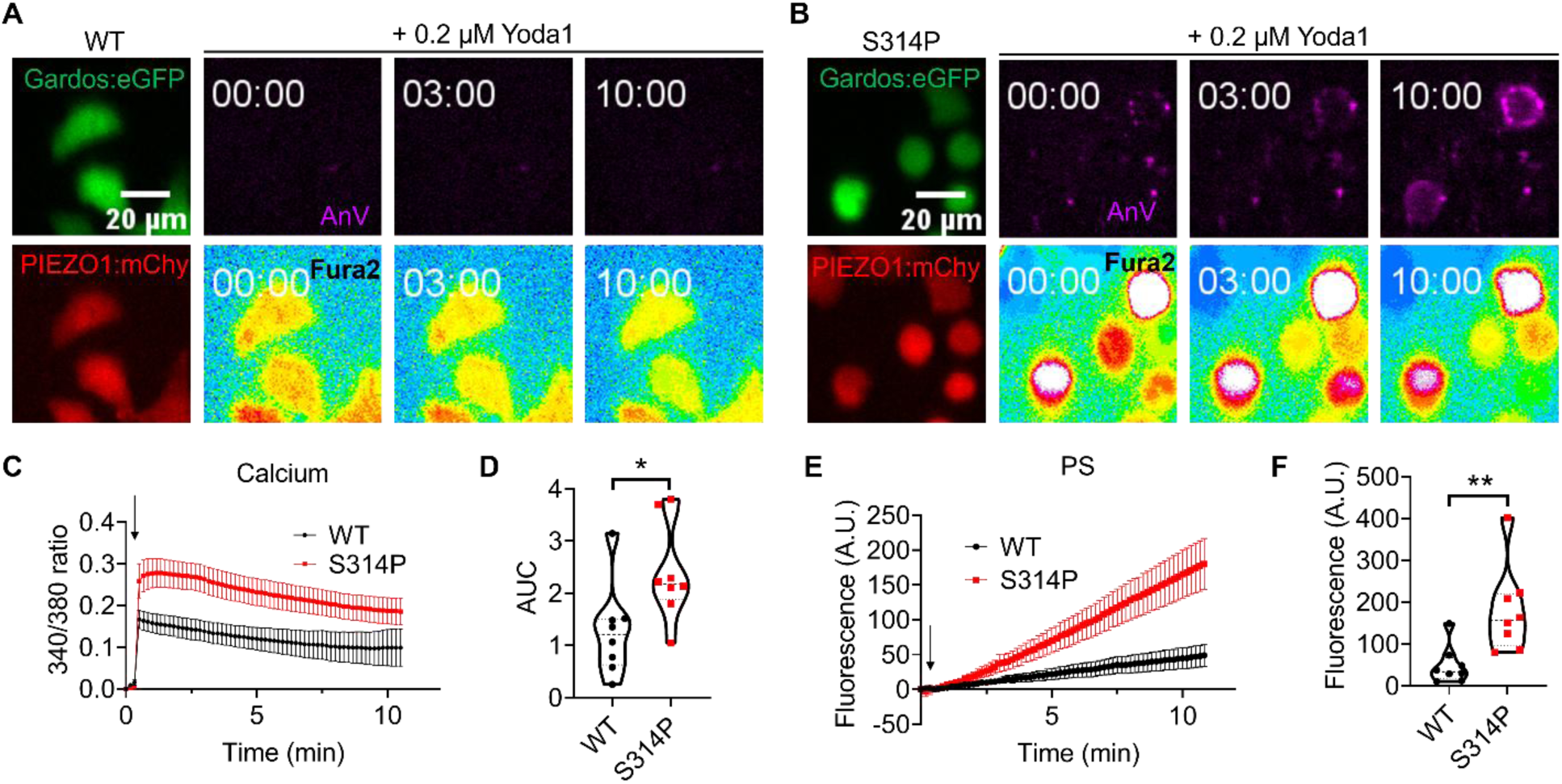
Gain-of-function Gardos variant further enhances PIEZO1-TMEM16F coupling in HEK293T cells. (A-B) Representative images of real-time fluorescence imaging of HEK293T PIEZO1 stable cells overexpressing WT Gardos (A) or Gardos S314P mutation (B). Cells expressing Gardos are represented in green, and cells expressing PIEZO1 are represented in red. PS exposure, as measured by AnV, is shown in magenta. Ca^2+^ intensity, as measured by Fura2, is shown in color gradients with blue indicating the lowest intensity, followed by green, yellow, red, and white. (C) Ca^2+^ influx measurement of HEK293T PIEZO1 stable cells overexpressing WT Gardos or Gardos S314P. The arrow indicates stimulation of 0.2 µM Yoda1. The error bars indicate ± SEM of 8 replicates in each condition. (D) AUC statistical analysis of Ca^2+^ influx results from panel C; two-sided t-test, * *P* = 0.0271 (n = 8). (E) PS exposure of HEK293T PIEZO1 stable cells overexpressing WT or S314P Gardos. (F) Statistical analysis of the fluorescence intensity of the last time points in panel E; two-sided t-test, ** *P* = 0.0055 (n = 8).

### Inhibition of Gardos attenuates PIEZO1-TMEM16F coupling in leukemia cells

The role of Gardos as an amplifier in PIEZO1-TMEM16F coupling suggests that genetic or pharmacological inhibition of Gardos may represent an unexplored strategy to reduce pathological PS exposure. We first tested this hypothesis in K562 cells, a human erythroleukemia cell line that recapitulates key properties of RBCs and is amenable to genetic manipulation^58,59^. Previous studies have demonstrated that both PIEZO1 and Gardos are expressed in K562 cells^60–63^. Our flow cytometry experiments showed that ionomycin induced robust lipid scrambling, which was absent in TMEM16F-deficient K562 cells, confirming functional expression of this CaPLSase in K562 cells (Fig. S1). Fluorescence imaging further revealed that stimulation of WT K562 cells with 0.5 µM Yoda1 elicited a rapid rise in intracellular Ca^2+^ followed by strong PS exposure (Fig. 5A, C-F), demonstrating strong PIEZO1-TMEM16F coupling in this model system.

**Figure 5.**
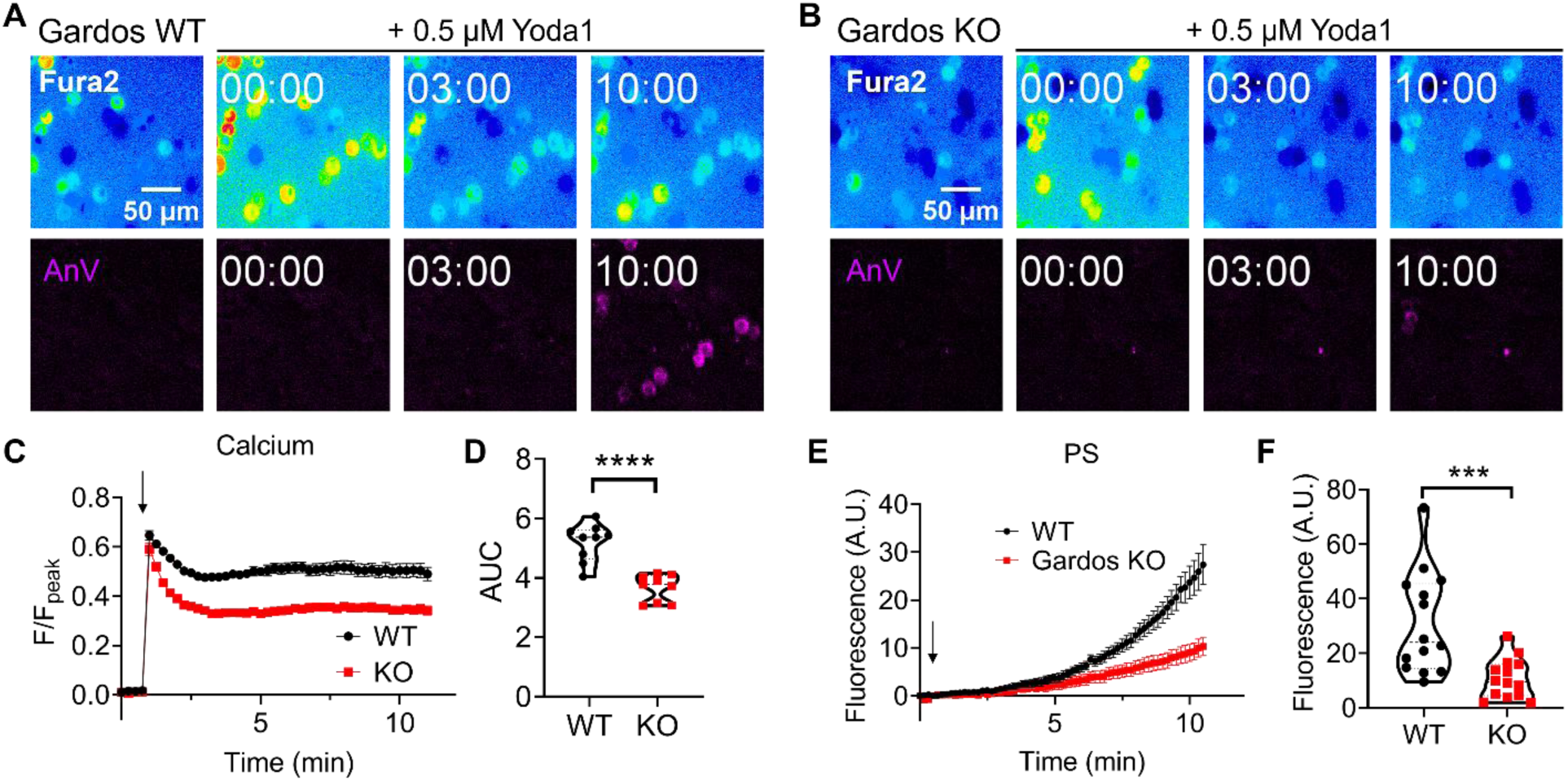
Gardos deficiency breaks PIEZO1-TMEM16F coupling in K562 cells. (A-B) Time-lapse images of Ca^2+^ levels indicated by Fura2 (top) and PS exposure indicated by fluorophore-conjugated AnV (magenta; bottom) of WT K562 cells (A) and Gardos knockout (KO) K562 cells (B). The intensity of Fura2 340/380 is shown in color gradients with blue indicating the lowest intensity, followed by green, yellow, red, and white. (C) Summary of Yoda1-induced Ca^2+^ influx of WT K562 cells versus Gardos KO K562 cells as measured by Fura2 ratiometric Ca^2+^ dye with a microplate reader. The 340/380 ratios were normalized by the respective peak values to account for differences in cell counts. The arrow indicates stimulation of 0.5 µM Yoda1. (D) The AUC statistical analysis of Ca^2+^ levels of WT versus Gardos KO K562 cells; **** two-sided t-test *P* < 0.0001 (n = 9). (E) Summary of Yoda1-induced PS exposure of WT K562 cells versus Gardos KO K562 cells. (F) End point fluorescence statistical analysis of PS exposure of WT versus Gardos KO K562 cells; *** two-sided t-test *P* = 0.0009 (n = 14).

To address the role of Gardos, we generated a *KCNN4* knockout (KO) K562 cell line. Whole-cell patch-clamp recordings in WT K562 cells detected a prominent inward K^+^ current that was sensitive to Senicapoc, a potent Gardos inhibitor (Fig. S2A, C-D). This Senicapoc-sensitive current was largely abolished in *KCNN4* KO cells (Fig. S2B-D), confirming successful genetic deletion of Gardos. Notably, compared with WT cells, *KCNN4* KO cells exhibited significantly reduced Yoda1-induced Ca^2+^ elevation and subsequent PS exposure (Fig. 5B-F). Our genetic approach thus demonstrated that Gardos also plays an essential regulatory role in PIEZO1-TMEM16F coupling in K562 cells.

We next tested whether pharmacological inhibition of Gardos produces similar effects. Current-clamp recording showed that Yoda1-induced strong membrane hyperpolarization was rapidly reversed by Senicapoc (Fig. S3A-B), indicating that Gardos activity underlies this response. Consistent with the loss of Gardos-mediated hyperpolarization (i.e., a reduced electrochemical driving force for Ca^2+^ entry), Senicapoc significantly attenuated PIEZO1-mediated Ca^2+^ influx (Fig. S3C-D). Flow cytometry analysis further demonstrated that Senicapoc decreased PS exposure in K562 cells (Fig. S3E-F). Taken together, these findings show that both genetic ablation and pharmacological inhibition of Gardos weaken PIEZO1-TMEM16F coupling and reduce Ca^2+^-dependent lipid scrambling, supporting the amplification role of Gardos in the PIEZO1-Gardos-TMEM16F axis.

### Common Gardos inhibitors exhibit off-target effects in RBCs

We next tested whether the commonly used Gardos inhibitors could attenuate PIEZO1-TMEM16F coupling and prevent PS exposure in healthy and diseased RBCs. We first focused on clotrimazole (CLT), a widely used Gardos inhibitor that was previously reported to reduce Ca^2+^-induced PS exposure in SCD RBCs^64^. Flow cytometry analysis showed that 20 µM CLT robustly inhibited Yoda1-induced PS exposure in HD, HX, and SCD RBCs (Fig. 6A-F). Our imaging and patch clamp recordings showed that CLT did not affect Ca^2+^-activated TMEM16F lipid scrambling (Fig. S4A-B) and currents (Fig. S4C-D) in HEK293T cells stably expressing TMEM16F, indicating that CLT at this concentration targets the PIEZO1-Gardos coupling without affecting scramblase activity.

**Figure 6.**
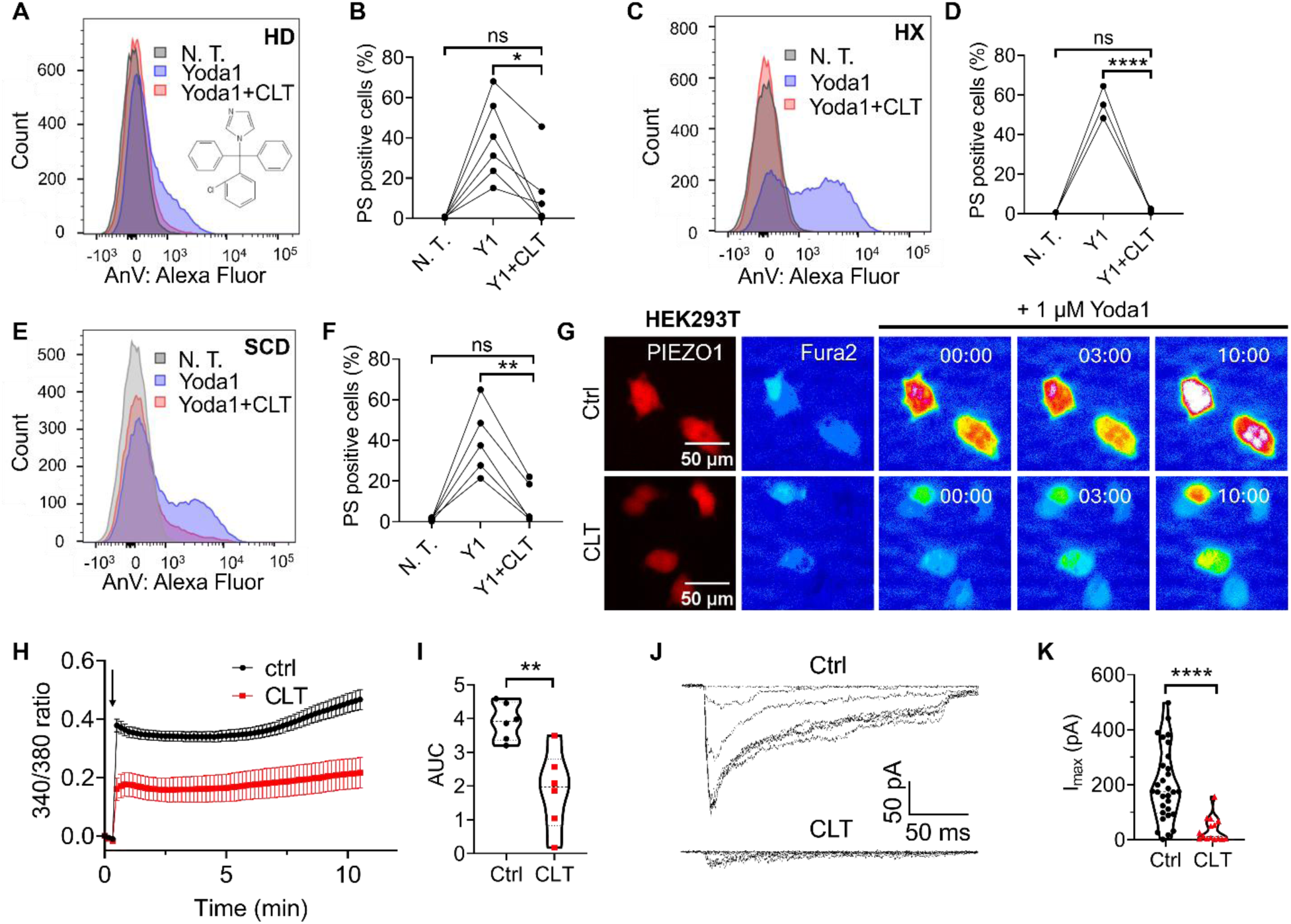
Clotrimazole inhibits PS exposure via PIEZO1 inhibition. (A) Representative flow cytometry traces of PS exposure of HD RBCs 10 minutes after 2 µM Yoda1 stimulation with or without 20 µM clotrimazole. The chemical structure of clotrimazole is included in the lower right corner. (B) Statistical summary of PS exposure of HD RBCs from six participants with or without clotrimazole; one-way ANOVA * *P* = 0.0291; ns *P* = 0.158 (n = 6). (C) Representative flow cytometry traces of PS exposure of HX RBCs 10 minutes after 2 µM Yoda1 stimulation with or without 20 µM clotrimazole. (D) Statistical summary of PS exposure of HX RBCs from one participation with or without clotrimazole; one-way ANOVA **** *P* < 0.0001; ns *P* = 0.3673 (n = 3 technical replicates). (E) Representative flow cytometry traces of PS exposure of SCD RBCs 10 minutes after 2 µM Yoda1 stimulation with or without 20 µM clotrimazole. (F) Statistical summary of PS exposure of SCD RBCs from five participants with or without clotrimazole; one-way ANOVA ** *P* = 0.009; ns *P* = 0.1236 (n = 5). (G) Time-lapse Ca^2+^ imaging of HEK293T cells stably expressing PIEZO1 (red). Cells were stimulated by 1 µM Yoda1 with or without the presence of 20 µM clotrimazole. Ca^2+^ levels were measured by the ratiometric Fura2 dye. (H) Summary of Yoda1-induced Ca^2+^ influx of HEK293T PIEZO1 stable cells with and without 20 µM clotrimazole. The arrow indicates 1 µM Yoda1 stimulation. (I) The AUC statistical analysis of HEK293T cells Ca^2+^ influx results from panel H; ** two-sided t-test *P* = 0.003 (n = 6). (J) Representative pressure-clamp traces of HEK293T PIEZO1 stable cells with and without 20 µM clotrimazole in the pipette solution. The pressure ranged from 0 to -60 mmHg with a holding potential at -80 mV under cell-attach configurations. (K) The maximum PIEZO1 current size with (n = 15) and without (n = 30) 20 µM clotrimazole solution; **** two-sided t-test *P* < 0.0001. N. T.: no treatment; Y1: Yoda1; CLT: clotrimazole.

Unexpectedly, we found 20 µM CLT also suppressed Yoda1-induced Ca^2+^ elevation in HEK293T cells stably expressing PIEZO1 (Fig. 6G-I). As there is no Gardos expression in this cell line, our results suggest that CLT may also target PIEZO1. To test this, we measured CLT’s effect on PIEZO1 channel activity using cell-attached pressure-clamp recordings. 20 µM CLT significantly inhibited force-induced PIEZO1 currents (Fig. 6J-K) in HEK293T cells stably expressing PIEZO1, confirming its direct inhibitory action on PIEZO1. These findings revealed a previously unrecognized off-target action of CLT on PIEZO1 and indicate that its suppression of PS exposure likely arises from concurrent inhibition of both Gardos and PIEZO1.

As CLT and TRAM-34, another widely used Gardos inhibitor, share structural similarity (Figs. 6A, S5A), we next tested whether TRAM-34 also inhibits PIEZO1. Ca^2+^ imaging showed that 20 µM TRAM-34 also strongly inhibited 1 µM Yoda1-induced Ca^2+^ elevation in PIEZO1 stable HEK293T cells (Fig. S5B-D). Consistently, pressure clamp recordings revealed near-complete inhibition of PIEZO1 currents, confirming that TRAM-34 also directly inhibits PIEZO1 (Fig. S5E-F).

Given the off-target effects of CLT and TRAM-34, we next evaluated Senicapoc, a Gardos inhibitor that advanced to phase III clinical trials for SCD^65^. In contrast to CLT and TRAM-34, Senicapoc did not alter PIEZO1-mediated Ca^2+^ entry (Fig. S6A-C) or channel activity (Fig. S6D-E), nor did it affect TMEM16F-dependent lipid scrambling (Fig. S7A-C) or currents in HEK293T cells (Fig. S7D-E). However, Senicapoc potentiated, rather than inhibited, Yoda1-induced PS exposure in HD (Fig. 7A-D), SCD (Fig. 7E-F), and HX (Fig. 7G-H) RBCs. This potentiation was evident at concentrations as low as 0.1 µM in HD RBCs (Fig. S8A-B) and was not attributable to hemolysis (Fig. S8C-D) or increased Ca^2+^ influx (Fig. S8E-F).

**Figure 7.**
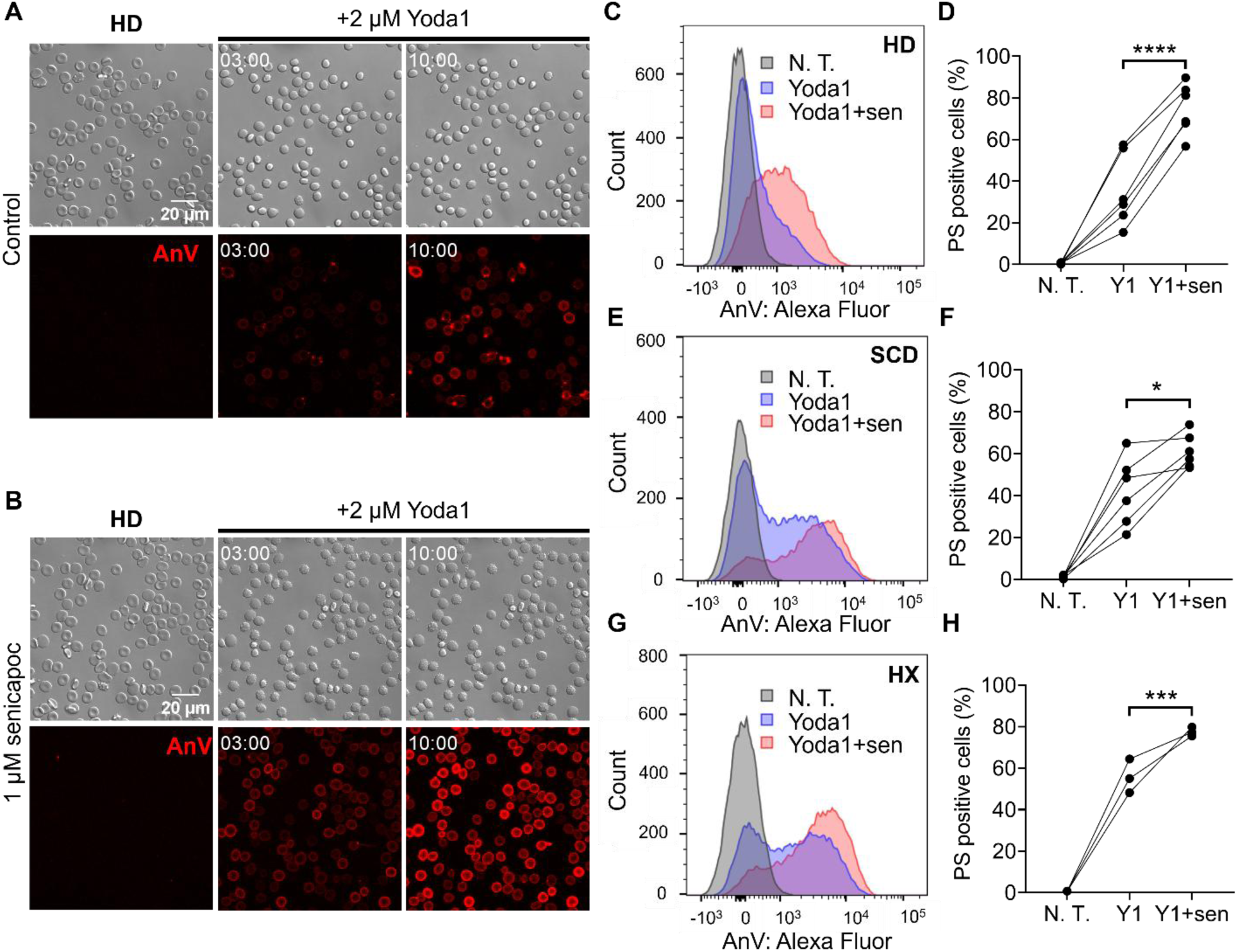
Senicapoc paradoxically promotes PS exposure in RBCs. (A-B) Representative confocal microscopy images of HD RBCs stimulated by 2 µM Yoda1 with (B) or without (A) 1 µM Senicapoc. (C) Representative flow cytometry traces of PS exposure of HD RBCs 10 minutes after 2 µM Yoda1 stimulation with or without 1 µM Senicapoc. (D) Statistical analysis of PS exposure of HD RBCs from six participants with or without Senicapoc. The connected dots indicate the same participant; **** two-sided t-test *P* < 0.0001 (n = 6). (E) Representative flow cytometry traces of PS exposure of SCD RBCs 10 minutes after 2 µM Yoda1 stimulation with or without 1 µM Senicapoc. (F) Statistical analysis of PS exposure of SCD RBCs from six participants with or without Senicapoc; * two-sided t-test, *P* = 0.0136 (n = 6). (G) Representative flow cytometry measurement of PS exposure of HX RBCs 10 minutes after 2 µM Yoda1 stimulation with or without 1 µM Senicapoc. (H) Statistical analysis of PS exposure of HX RBCs from one participant with or without Senicapoc. The connected dots indicate technical replicates; *** two-sided t-test, *P* = 0.0006 (n = 3; technical replicates). N. T.: no treatment; Y1: Yoda1; sen: Senicapoc.

Collectively, these results demonstrate that although Gardos is a key amplifier of PIEZO1-TMEM16F coupling, commonly used Gardos inhibitors, including CLT, TRAM-34, and Senicapoc, exhibit off-target or unexpected RBC-specific effects that preclude their straightforward application in suppressing this pathway in RBCs. These findings underscore the need to develop more selective Gardos inhibitors to therapeutically modulate pathological PS exposure.

## DISCUSSION

The Gardos channel has long been recognized as a key regulator of RBC ion and volume homeostasis and has been implicated in various RBC disorders^37,49,66–69^. Traditionally, Gardos was viewed primarily as a Ca²⁺ effector downstream of PIEZO1 that mediates K⁺ efflux, thereby promoting membrane hyperpolarization, RBC dehydration, and volume regulation^38,44–47^. Our study expands this view by identifying Gardos as a feedback regulator of RBC Ca²⁺ signaling that is critical for efficient TMEM16F activation. We show that PIEZO1-mediated Ca²⁺ entry activates Gardos, whose opening hyperpolarizes the RBC membrane and increases the electrochemical driving force for additional Ca²⁺ influx through PIEZO1. This positive-feedback mechanism amplifies sustained Ca²⁺ signaling, promotes TMEM16F Ca²⁺-activated phospholipid scramblase activity, and drives PS exposure. Thus, Gardos is not simply a passive effector of Ca²⁺ signaling, but a central amplifier that actively links mechanosensitive Ca²⁺ entry to lipid scrambling and pathological membrane remodeling.

This mechanism refines the current model of PIEZO1-TMEM16F coupling in RBCs^11^. A simple two-component model, in which PIEZO1 directly supplies the Ca²⁺ required for TMEM16F activation, does not fully explain how limited PIEZO1 activity generates sufficient Ca²⁺ elevation to trigger robust TMEM16F activation and PS exposure. This issue is particularly relevant in mature RBCs, where PIEZO1 is present at low copy number and exhibits rapid inactivation^29–31^, whereas TMEM16F requires relatively high Ca²⁺ concentrations for efficient activation^22,32–36^. Gardos-dependent membrane hyperpolarization provides a unique solution to this problem by increasing the driving force for Ca²⁺ entry through PIEZO1. More broadly, the ability of Gardos to increase the electrochemical driving force for Ca²⁺ entry is unlikely to be limited to PIEZO1-Gardos coupling. In principle, Gardos activation could amplify Ca²⁺ influx through any Ca²⁺ permeable channel, provided that the initial Ca²⁺ entry is sufficient to activate Gardos. RBCs have been reported to express additional Ca²⁺ permeable pathways, including TRPV2 and NMDA receptors^70–72^. Whether Gardos functionally couples to these channels to enhance Ca^2+^ influx and promote TMEM16F-dependent phospholipid scrambling remains an important question for future studies. Additionally, the observation of this mechanism in HEK293T cells and erythroleukemia cells demonstrates that the PIEZO1-Gardos-TMEM16F axis identified in this study can exist in cell types beyond mature RBCs, where this functional module transduces mechanical stimulation into dynamic phospholipid scrambling and membrane remodeling.

Gardos amplification has important implications for RBC disorders characterized by abnormal cation flux, dehydration, and PS exposure. In PIEZO1 GOF HX RBCs, enhanced PIEZO1 activity is expected to increase Ca²⁺ entry and activate Gardos, thereby further amplifying Ca²⁺ influx and promoting TMEM16F-mediated PS exposure^73–80^. Consistent with this model, the Gardos channelopathy variant S314P^81^ enhanced PIEZO1-mediated Ca²⁺ influx and PS exposure in our reconstituted system, suggesting that a similar feed-forward amplification mechanism may operate in RBCs from patients with GOF *KCNN4* channelopathy. Future studies examining PS exposure in Gardos channelopathy RBCs will be important for determining whether this mechanism contributes to disease pathophysiology.

This model is also relevant to SCD. Gardos has long been considered a therapeutic target in SCD because K⁺ and water loss increase intracellular hemoglobin concentration^65,67,69,82–85^, thereby promoting HbS polymerization, sickling, and further dehydration of RBCs. Our results suggest an additional mechanism by which Gardos may contribute to SCD pathophysiology: amplification of PIEZO1-mediated Ca²⁺ influx and TMEM16F-dependent PS exposure. Because PS-positive SCD RBCs and RBC-derived microparticles contribute to thrombin generation, endothelial adhesion, vascular inflammation, and vaso-occlusive complications^15,16,20,21,86–90^, Gardos inhibition could, in principle, provide dual benefits by reducing both RBC dehydration and procoagulant membrane remodeling. However, SCD RBCs are affected by multiple overlapping stress pathways, including HbS polymerization, oxidative damage, altered metabolism, membrane fragility, and dysregulated ion transport^91,92^. Therefore, Gardos amplification should be viewed as one important contributor to pathological PS exposure rather than the sole driver.

Our pharmacological findings highlight both the promise and the challenge of targeting Gardos. CLT and TRAM-34, two commonly used Gardos inhibitors, strongly suppressed Yoda1-induced Ca²⁺ influx and PS exposure, but both compounds also directly inhibited PIEZO1 at micromolar concentrations. Therefore, their effects on PIEZO1-TMEM16F coupling cannot be attributed solely to Gardos inhibition. Dual inhibition of PIEZO1 and Gardos could be therapeutically attractive in principle, especially in diseases such as SCD where both Ca²⁺ influx and RBC dehydration contribute to pathology. However, the adverse effects of CLT may limit its long-term therapeutic use^83,93^.

Senicapoc revealed a distinct difference. Unlike CLT and TRAM-34, Senicapoc did not detectably affect PIEZO1 or TMEM16F activity in our assays. Unexpectedly, however, Senicapoc potentiates Yoda1-induced PS exposure in HD, SCD, and HX RBCs at concentrations as low as 0.1 µM. This effect was not attributable to hemolysis or increased Ca²⁺ influx, suggesting an unknown RBC-specific mechanism that is absent in HEK293T or K562 cells. Because previously reported steady-state plasma concentrations of Senicapoc are in a similar range^65,67^, this effect should be carefully considered in future studies of Senicapoc and related Gardos inhibitors. The phase III clinical trial of Senicapoc showed that, despite ameliorated RBC dehydration, lessened hemolysis, and increased RBC survival, Senicapoc increased the frequency of vaso-occlusion in SCD patients^65^. As PS exposure on RBCs enhances adhesion towards endothelial cells^10^, the Senicapoc effect of enhancing PS exposure may contribute to increased vaso-occlusion in the clinical trial. Further work is needed to define the molecular basis of Senicapoc-potentiated PS exposure and assess whether this effect is relevant to its clinical performance in SCD.

In summary, this study identifies Gardos as a critical amplifier of PIEZO1-TMEM16F coupling in RBCs. By linking mechanosensitive Ca²⁺ entry to membrane hyperpolarization and TMEM16F-dependent PS exposure, Gardos connects RBC ion homeostasis, dehydration, and procoagulant membrane remodeling. These findings provide a mechanistic framework for understanding excessive PS exposure in SCD, PIEZO1-HX, and Gardos channelopathy, while also revealing important limitations of commonly used Gardos inhibitors. Future studies should determine whether Gardos-amplified lipid scrambling contributes to the pathophysiology of other blood disorders and should prioritize the development of selective Gardos modulators that mitigate pathological RBC signaling with improved selectivity and safety.

## DATA AVAILABILITY

All study data are included in the article and/or SI Appendix. Contact Dr. Huanghe Yang for data and material requests at.

## ACKNOWLEDGEMENT

We thank Drs. Augustus J. Lowry and Ke Shan for constructing the HEK293T PIEZO1 stable cells. We thank Dr. Pengfei Liang for the intellectual input and technical assistance. We thank Dr. Miao Zhang for generously sharing the WT and GOF *KCNN4* plasmids. We also greatly appreciate SCD and HX participants, as well as healthy donors, for their blood sample donation. This work was supported by NIH-R35GM153196 (to HY), and the American Society of Hematology (ASH) Graduate Hematology Award (to YCSW, 2023–2025).

## AUTHORSHIP

Contributions: Conceptualization: HY; Methodology: YCSW, HY; Investigation: YCSW, MD; Data curation and analysis: YCSW; Visualization: YCSW, HY; Validation: YCSW; Writing – original draft: HY, YCSW; Writing – review & editing: all authors; Resources: HY, GL, SK, GMA, MJT; Supervision and project administration: HY; Funding acquisition: HY, YCSW.

## CONFLICT-OF-INTEREST DISCLOSURE

The authors declare no competing financial interests.

## Notes

### Competing Interest Statement

The authors have declared no competing interest.

