## Supplementary Methods and Figures for "Gardos potassium channel amplifies PIEZO1-TMEM16F coupling in red blood cells"

**The file includes:**

Supplementary Methods
Supplementary Figure 1 to Figure 8

### SUPPLEMENTARY METHODS

#### Reagents

Yoda1 (Cat# 21904), Senicapoc (Cat# 29679), Clotrimazole (Cat# 15278), TRAM-34 (Cat# 23385), and Fura2-AM (Cat# 14591) were purchased from Cayman Chemical Company (Ann Arbor, MI). Calbryte 520 (Cat# 20651) was purchased from AAT Bioquest (Pleasanton, CA). Poly-l-lysine (PLL) (Cat# P2636) and fetal bovine serum (FBS; Cat# F2442) were purchased from Sigma-Aldrich (St. Louis, MO). Annexin V's (Cat# 29005, Cat# 29011, and Cat#29014) were purchased from Biotium (Fremont, CA). Lipofectamine 2000 (Cat# 11668019), penicillin-streptomycin (Cat# 15140122), Hank's Balanced Salt Solution (HBSS Cat# 14025-092), Dulbecco's Phosphate Buffered Saline (DPBS; Cat# 14190-144), Iscove's Modified Dulbecco's Medium (IMDM; Cat# 12440-053), Dulbecco's Modified Eagle Medium (DMEM; Cat# 11995-065), and $Ca^{2+}$ free DMEM (Cat# 21068-028) were purchased from Thermo Fisher Scientific (Waltham, MA). The WT and GOF Gardos-IRES-eGFP plasmids were kind gifts from Dr. Miao Zhang.

#### Cell culture and transfection

The K562 human erythroleukemia cell line was purchased from and authenticated by the Duke Life Science Facility. The Gardos KO (sgRNA: TGGCATGAAAGGCCACGATG) and TMEM16F KO (sgRNA: AAAAGTACACGCACCATGGG) K562 cell line were generated and sequence confirmed by the Duke Functional Genomics Core. The HEK293T cell line stably expressing eGFP-tagged TMEM16F was a kind gift from Drs. Min Li and Lily Jan as described previously[11]. The HEK293T cell line stably expressing hPIEZO1-IRES-mCherry was generated using lentivirus. Briefly, the lentiviral construct (derived from Addgene plasmid #62554) was modified to replace the puromycin cassette with a hygromycin resistance gene (Addgene #60498) and then the human PIEZO1 cDNA (GenBank AGH27891.1) was inserted downstream of an IRES. Empty vector control lacked the PIEZO1-IRES segment, while having the mCherry segment. Constructs were assembled using In-Fusion Snap Assembly (Takara, #638947). Lentivirus was produced in HEK293T cells by co-transfecting packaging plasmids (pMD2.G and psPAX2, Addgene #12259 and #12260) with Lipofectamine 2000. WT HEK293T cells were transduced by the lentiviruses in the presence of 10 μg/ml polybrene and selected using hygromycin.

Cells were cultured in IMDM and DMEM, supplemented with 10% FBS and 1% penicillin/streptomycin, for K562 and HEK293T cells, respectively. All cells were cultured in a humidified atmosphere with 5% CO2 at 37°C.

To perform transfection, HEK293T cells were seeded on 0.1 mg/mL PLL-coated coverslips and transfected with Lipofectamine 2000 mixed with the appropriate plasmids according to the manufacturer's guidelines. The DMEM media containing the transfection reagents were changed to $Ca^{2+}$-free DMEM after 4 hours. Experiments were performed 24 to 48 hours after transfection.

**Electrophysiology**

Patch clamp recordings were performed using an Axopatch 200B amplifier (Molecular Devices, Inc., San Jose, CA). Signals were low-pass filtered at 5 kHz, digitized at 10 kHz with a Digidata 1550A digitizer, and acquired using Clampex 10 software. Data were analyzed using Clampfit 11.4.3 software (Molecular Devices, Inc., San Jose, CA). Glass pipettes were pulled from borosilicate capillaries (Sutter Instruments, Novato, CA) and subsequently fire-polished using a microforge (Narishige, Amityville, NY) to obtain a final resistance of 1.7–3.0 MΩ.

For current clamp measurement of Gardos-mediated membrane hyperpolarization, the bath solution contained (in mM) 140 NaCl, 5 KCl, 10 HEPES, and 5 EGTA at pH 7.3; the pipette solution contained (in mM) 140 KCl, 1 $MgCl_2$, 10 HEPES, and 0.2 EGTA at pH 7.4; the $Ca^{2+}$-

solution for bath perfusion upon whole-cell configuration formation contained (in mM) 140 NaCl, 5 KCl, 2 $MgCl_2$, 10 HEPES, and 2 $CaCl_2$ at pH 7.4. Yoda1 was pre-diluted in the $Ca^{2+}$-perfusion solution. Whole-cell patch configuration was first formed in the $Ca^{2+}$-free bath solution. Once a stable whole-cell patch was formed, the membrane potential was measured under the current clamp. Yoda1 was then perfused, using a valve-controlled manifold (ALA-VM8, ALA Scientific Instruments), to the cells while the membrane potential was continuously recorded.

For TMEM16F current measurement, HEK293T cells stably expressing mTMEM16F were seeded on 0.1 mg/mL PLL-coated coverslips overnight. Whole-cell voltage-clamp was performed using the bath solution (in mM): 150 NaCl and 10 HEPES at pH 7.4; the pipette solution (in mM): 150 NaCl, 1 $CaCl_2$, and 10 HEPES at pH 7.4. Voltage steps ranged from -100 to +120 mV in +20 mV increments. Holding potential was -60 mV. Senicapoc or Clotrimazole was diluted in the bath solution and perfused to the cells.

For the measurement of PIEZO1 current, HEK293T cells stably expressing PIEZO1 were seeded on 0.1 mg/mL PLL-coated coverslips overnight. The cell-attached pressure clamp (ALA Scientific Instruments, model HSPC-2-SB) was used to measure PIEZO1 current at a holding potential of -80 mV. The pressure steps ranged from 0 to -60 mmHg in -10 mmHg increments. The bath solution contained (in mM) 150 NaCl, and 10 HEPES at pH 7.4, and the pipette solution was the bath solution with an additional 2 mM $MgCl_2$. Senicapoc or Clotrimazole was included in the pipette solution as needed.

For measuring the Senicapoc-sensitive currents of K562 WT and Gardos KO cells, we used whole-cell voltage clamp. The bath solution contained (in mM) 140 KCl, 10 HEPES, and 5 EGTA at pH 7.3; the pipette solution contained (in mM) 140 KCl, 10 HEPES, 5 EGTA, and ~5 µM free $Ca^{2+}$

(estimated by WEBMAXC online software[48]). 20 µM Senicapoc was diluted in the bath solution and perfused to the cells as needed. Inward currents were measured at -100 mV.

**Microplate reader $Ca^{2+}$ measurement of K562 cells and RBCs**

For K562 cell $Ca^{2+}$ measurement, the cells were first collected and pelleted with centrifugation at 500 x g for 5 min. The packed cells were then re-diluted to $5 \times 10^6$ cells/mL in HBSS and kept on ice. 50 µL were diluted to 500 µL in HBSS containing 2 µM Fura2. The cells were incubated at 37°C for 45 minutes. The cells were centrifuged again to remove the Fura2 solution and resuspended in 500 µL HBSS. 50 µL of the cells were added to each well of the plate. Yoda1 solution was added to reach a final concentration of 0.5 µM. Fluorescence signal was measured with a plate reader in real-time, and the 340/380 emission ratio was calculated as an indicator of $Ca^{2+}$ levels.

For RBC $Ca^{2+}$ measurement, 10 µL of packed RBCs were diluted in 500 µL of HBSS containing 1 µM Calbryte 520. The cells were incubated at 37°C for 15 minutes. After Calbryte loading, the cells were centrifuged at 100 x g for 5 minutes to remove the Calbryte solution. The cell pellet was then resuspended in 500 µL HBSS. Senicapoc was added to the desired concentrations. 100 µL of RBC solution was added to the corresponding well of a 96-well plate in triplicate. Yoda1 solution was added to reach the desired final concentration. Fluorescence signal was measured with a plate reader in real-time with 490 nm excitation and 525 nm emission.

**Microplate reader measurement of RBC hemolysis**

10 µL of packed RBCs were diluted in 500 µL of HBSS in a 1.5 mL tube in triplicate for each condition. Senicapoc was added to the desired concentrations. The cells were then stimulated with Yoda1 at the desired concentration for 10 minutes at room temperature. 10 µL of packed RBCs were diluted in 500 µL of water in a 1.5 mL tube as a positive control for 100% hemolysis. The

cells were centrifuged at 1000 x g for 1 minute. 200 μL of supernatant was added to the corresponding well of a 96-well plate. Absorbance was measured at 577 nm with 690 nm for background subtraction.

**SUPPLEMENTARY FIGURES**

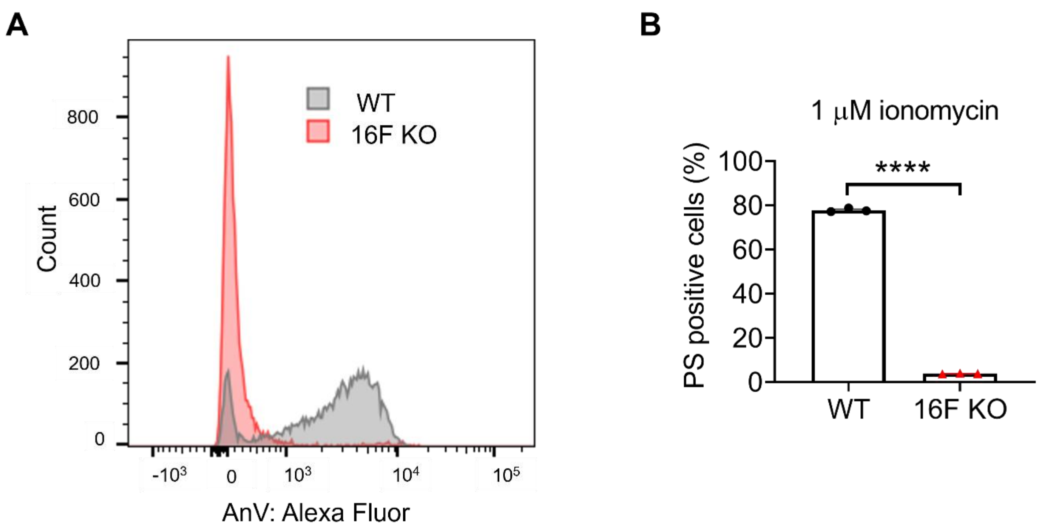


**Figure S1. TMEM16F is responsible for $Ca^{2+}$-induced PS exposure in K562 cells.** (A) Representative flow cytometry traces of wildtype (WT) and TMEM16F knockout (KO) K562 cells measuring PS exposure using AnV after 10 minutes of 1 μM ionomycin stimulation. (B) Statistical analysis of PS-exposed WT and TMEM16F KO K562 cells. Two-sided t-test ****, $P < 0.0001$ (n = 3).

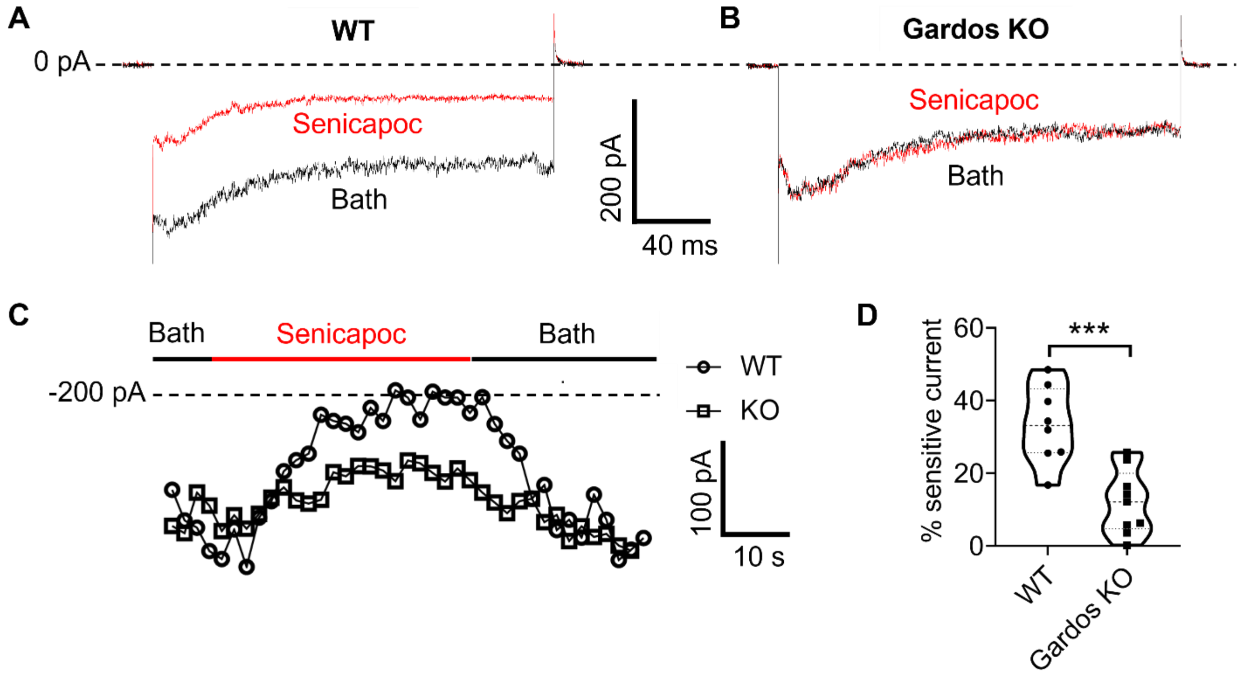


**Figure S2. Validation of K562 KCNN4/Gardos KO cells.** (A-B) Representative current traces of WT (A) and Gardos KO (B) K562 cells recorded using whole-cell voltage-clamp. The patches were held at 0 mV, and currents were elicited by a -100 mV voltage step. The red traces represent currents after 20 µM Senicapoc perfusion, while the black traces represent currents under bath solution perfusion. (C) Representative maximum currents over time for WT (circle) and Gardos (square) KO K562 cells. After the formation of a patch, the patch was perfused with the bath solution first (black), followed by 20 µM Senicapoc (red), and bath solution (black). (D) Percentage of Senicapoc-sensitive current calculated by ($I_{bath}$ – $I_{sen}$)/$I_{bath}$ x 100; two-sided t-test, *** $P$ = 0.0004 (n = 8, 9).

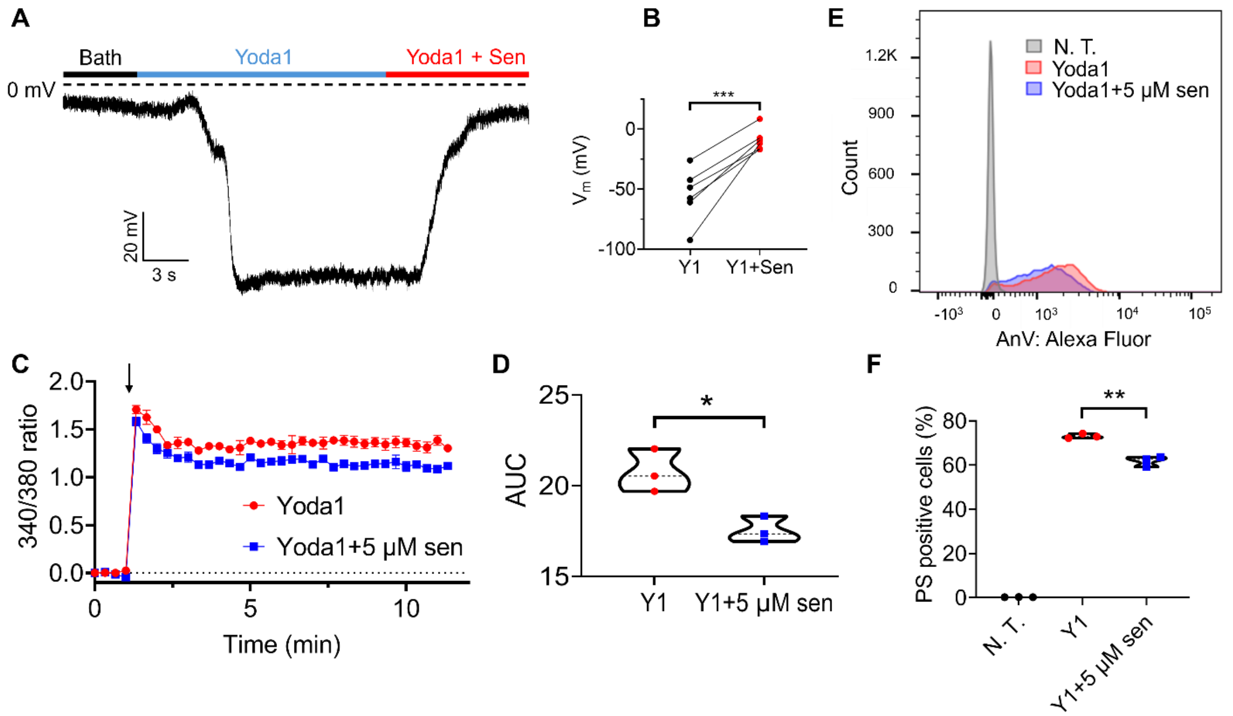


**Figure S3. Gardos antagonist Senicapoc breaks PIEZO1-TMEM16F coupling in K562 cells.** (A) A representative current clamp trace of the K562 cell membrane potential. After formation of the whole-cell configuration, 1 μM Yoda1 was perfused to the cells (cyan), followed by perfusion of 1 μM Yoda1+ 20 μM Senicapoc (red). (B) Statistical analysis summary of the K562 cell membrane potential during Yoda1 and Yoda1+Senicapoc perfusion. Connected dots indicate the same cell; two-sided t-test, *** $P = 0.0009$ (n = 6). (C) $Ca^{2+}$ levels of K562 cells were monitored by Fura2 ratiometric $Ca^{2+}$ dye using a microplate reader. The arrow indicates stimulation of 0.5 μM Yoda1. The error bars indicate ± SEM. (D) The AUC statistical analysis of $Ca^{2+}$ influx levels from panel C; two-sided t-test, * $P = 0.016$ (n = 3). (E) Representative flow cytometry traces of PS exposure as measured by AnV with or without Senicapoc after 10 minutes of 0.5 μM Yoda1 stimulation. (F) Statistical analysis of K562 cells PS exposure after 0.5 μM Yoda1 stimulation with or without Senicapoc; two-sided t-test, ** $P = 0.0016$ (n = 3). Y1: Yoda1; Sen: Senicapoc; N. T.: no treatment.

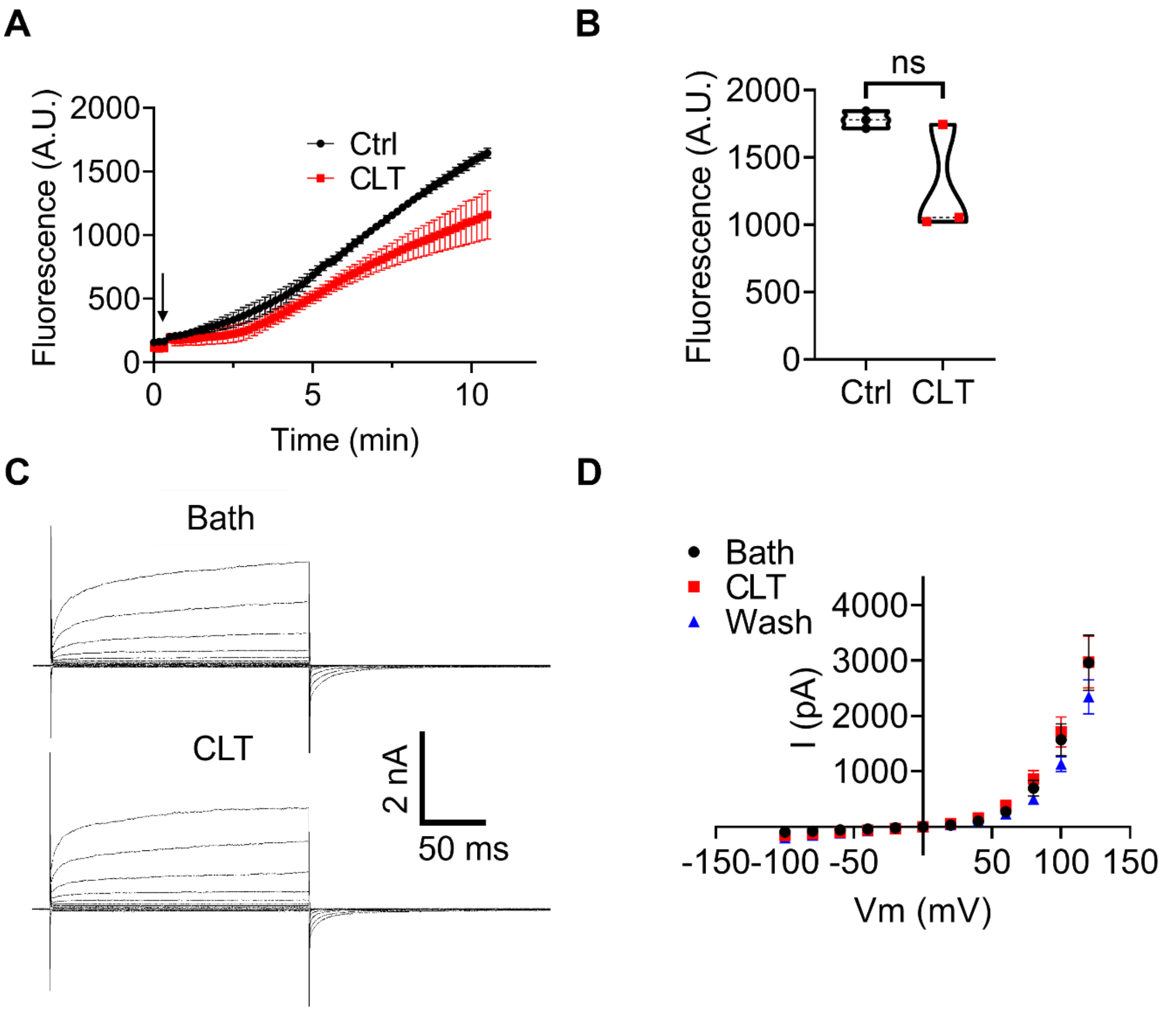


**Figure S4. Clotrimazole (CLT) does not affect TMEM16F activity.** (A) PS exposure measurement of HEK293T cells stably expressing mTMEM16F with (red) or without (black) 20 µM CLT. The arrow indicates stimulation with 1 µM ionomycin. The error bars represent ± SEM (n = 3). (B) Statistical analysis of endpoint fluorescence from panel A; two-sided t-test, ns (not significant) $P = 0.1002$ (n = 3). (C) Representative mTMEM16F current traces with or without 20 µM CLT measured by the whole-cell voltage clamp configuration. The two traces represent TMEM16F currents of the same cell under perfusion of bath solution or 20 µM CLT solution. (D) I-V relationship of TMEM16F currents under perfusion of bath (black), 20 µM CLT (red), and a second round of bath solution (wash; blue). The error bars represent ± SEM (n = 5).

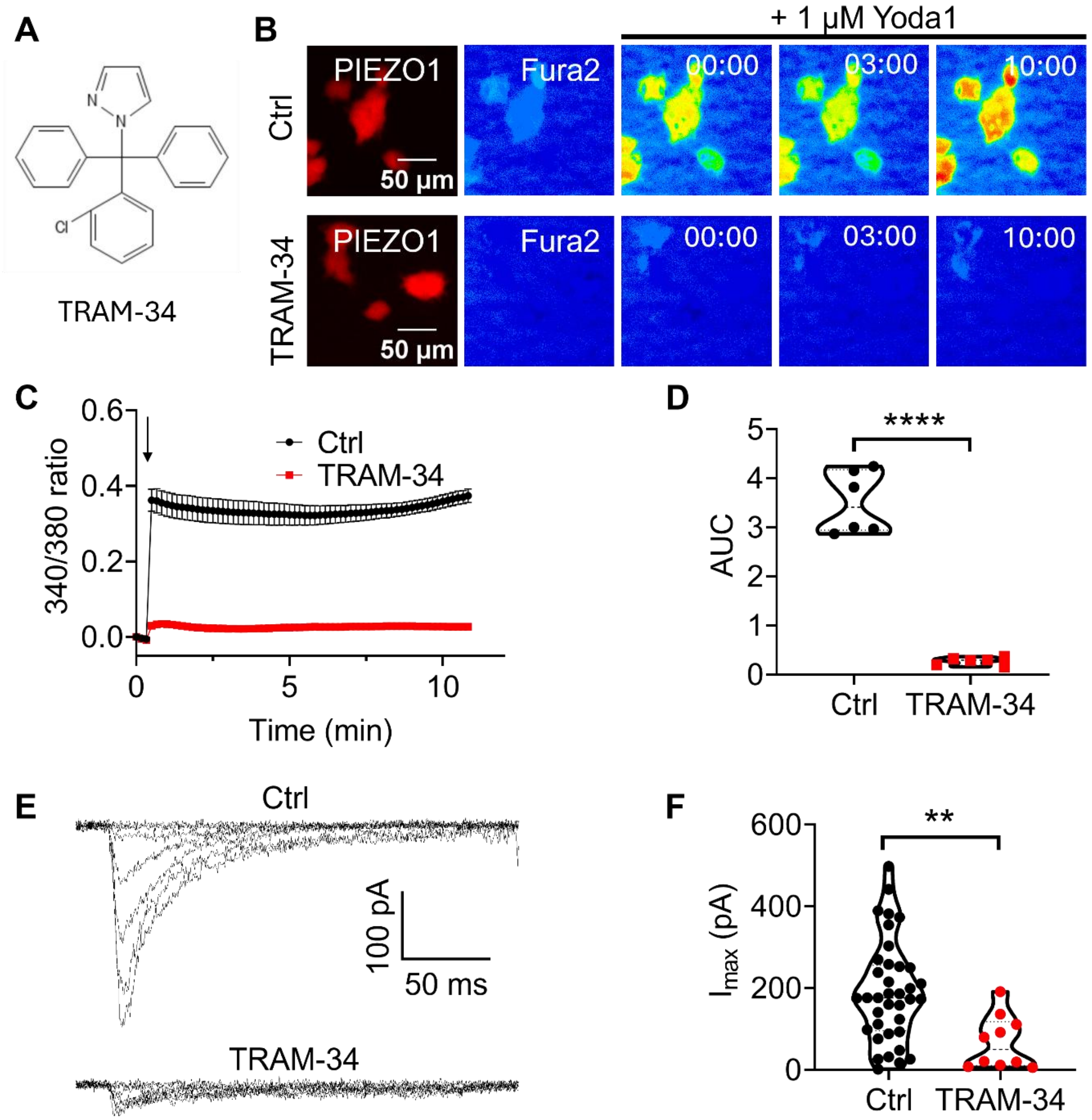


**Figure S5. TRAM-34 inhibits PIEZO1.** (A) The chemical structure of TRAM-34. (B) Representative Fura2 $Ca^{2+}$ imaging of HEK293T cells stably expressing PIEZO1 (red). $Ca^{2+}$ intensity is shown in color gradients with blue indicating the lowest intensity, followed by green, yellow, red, and white. Cells were stimulated with 1 μM Yoda1 with or without the presence of 20 μM TRAM-34. (C) Summary of $Ca^{2+}$ influx of HEK293T cells stably expressing PIEZO1 with (red) or without (black) the presence of 20 μM TRAM-34. The arrow indicates stimulation of 1 μM Yoda1. The error bars represent ± SEM (n = 6). (D) The AUC statistical analysis of $Ca^{2+}$ influx results from panel C; two-sided t-test, **** *P* < 0.0001. (E) Representative PIEZO1 current measured by cell-attached pressure-clamp with or without 20 μM of TRAM-34 in the pipette solution. The pressure ranged from 0 to -60 mmHg with a holding potential at -80 mV. (F) The maximum PIEZO1 current size with (n = 10) and without (n = 37) TRAM-34; two-sided t-test, ***P* = 0.0039.

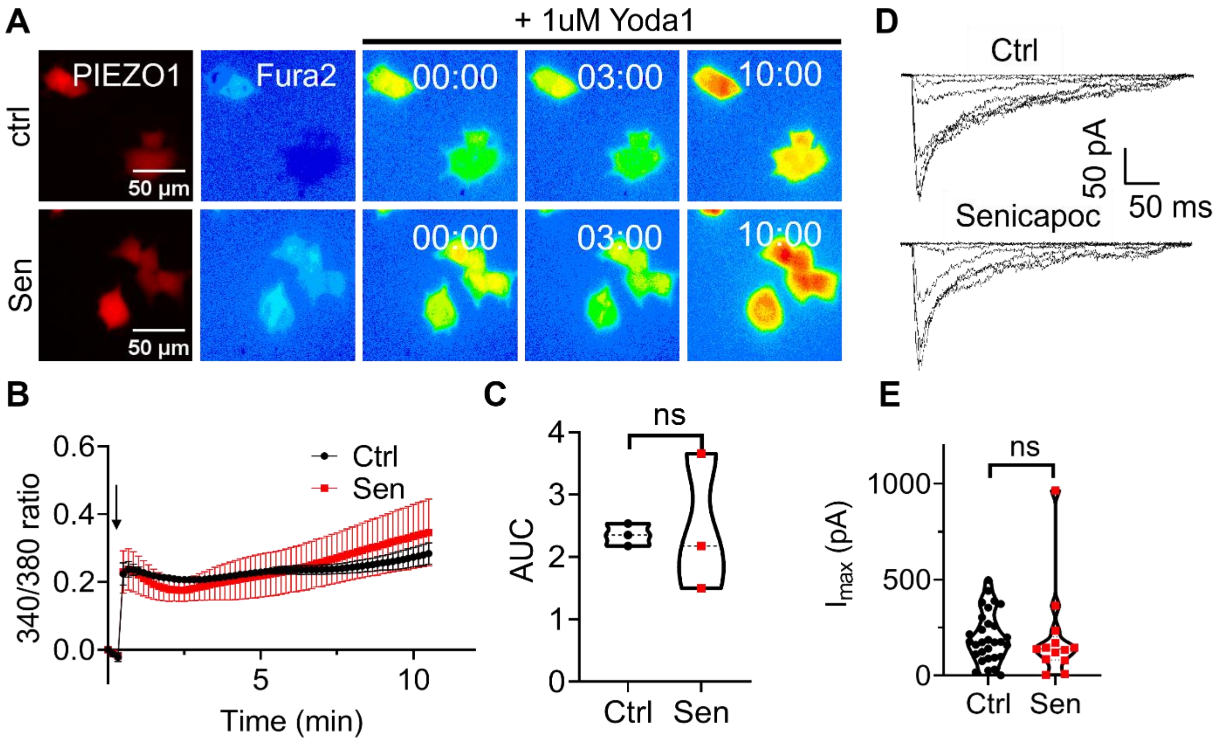


**Figure S6: Senicapoc does not affect PIEZO1.** (A) Representative Fura2 $Ca^{2+}$ images of HEK293T cells stably expressing PIEZO1 (red). Cells were stimulated with 1 µM Yoda1 with or without 20 µM Senicapoc. (B) Summary of $Ca^{2+}$ influx of HEK293T PIEZO1 stable cells with (red) or without (black) 20 µM Senicapoc as measured by real-time fluorescence imaging. The arrow indicates stimulation of 1 µM Yoda1. (C) The AUC analysis of $Ca^{2+}$ influx of HEK293T PIEZO1 stable cells with or without 20 µM Senicapoc from panel B; two-sided t-test, ns (not significant), *P* = 0.8955 (n = 3). (D) Representative cell-attached pressure-clamp trace of HEK293T PIEZO1 stable cells with and without 20 µM Senicapoc in the pipette solution. The pressure ranged from 0 to -60 mmHg with a holding potential at -80 mV. (E) Statistical analysis of peak PIEZO1 current size with (n = 13) or without 20 µM Senicapoc (n = 30); two-sided t-test, ns, *P* = 0.7278. Sen: Senicapoc.

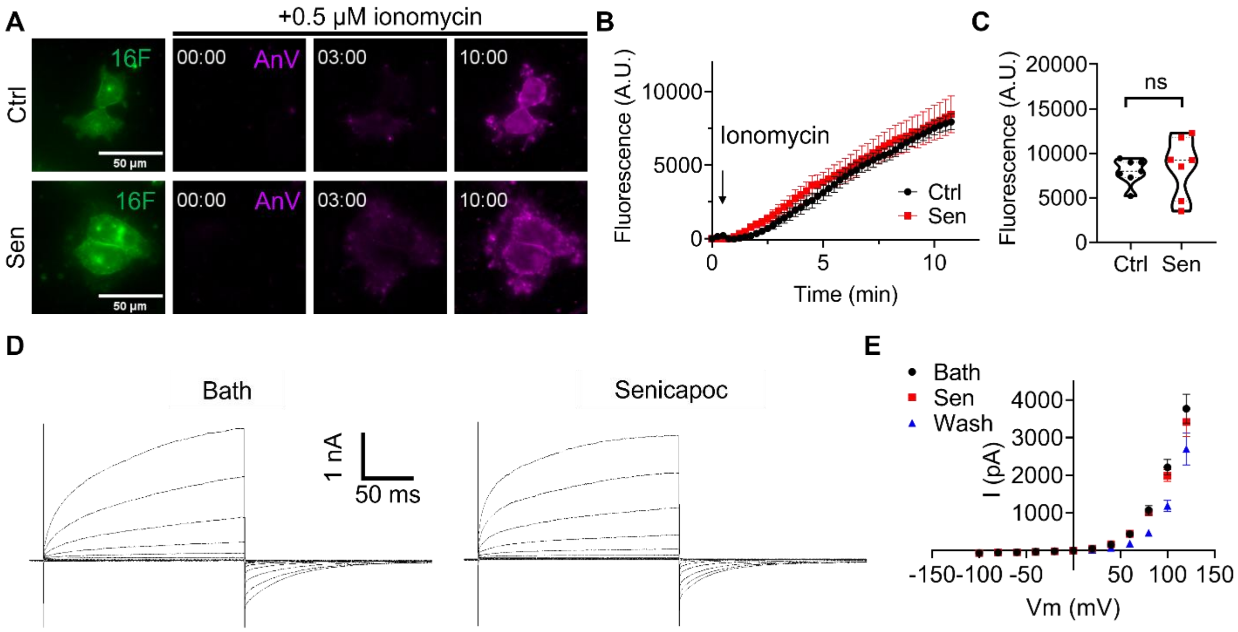


**Figure S7. Senicapoc does not affect TMEM16F activity.** (A) Representative time-lapse fluorescence images of PS exposure in mTMEM16F stable HEK293T cells with (bottom) or without (top) 20 µM Senicapoc in response to 0.5 µM ionomycin stimulation. Green represents mTMEM16F-eGFP expression, and magenta represents AnV binding to exposed PS. (B) Time course of PS exposure of mTMEM16F stable HEK293T cells with (red) or without (black) 20 µM Senicapoc. The arrow indicates stimulation of 0.5 µM ionomycin. (C) Statistical analysis of endpoint AnV fluorescence after 10 minutes of 0.5 µM ionomycin stimulation from panel B; two-sided t-test, ns (not significant), $P$ = 0.7111 (n = 7). (D) Representative whole-cell TMEM16F current traces before (bath) and after 20 µM Senicapoc from the same cell. (E) I-V relationship of TMEM16F currents under perfusion of bath, 20 µM Senicapoc, and a second round of bath solution (wash). The error bars represent ± SEM (n = 5). Sen: Senicapoc.

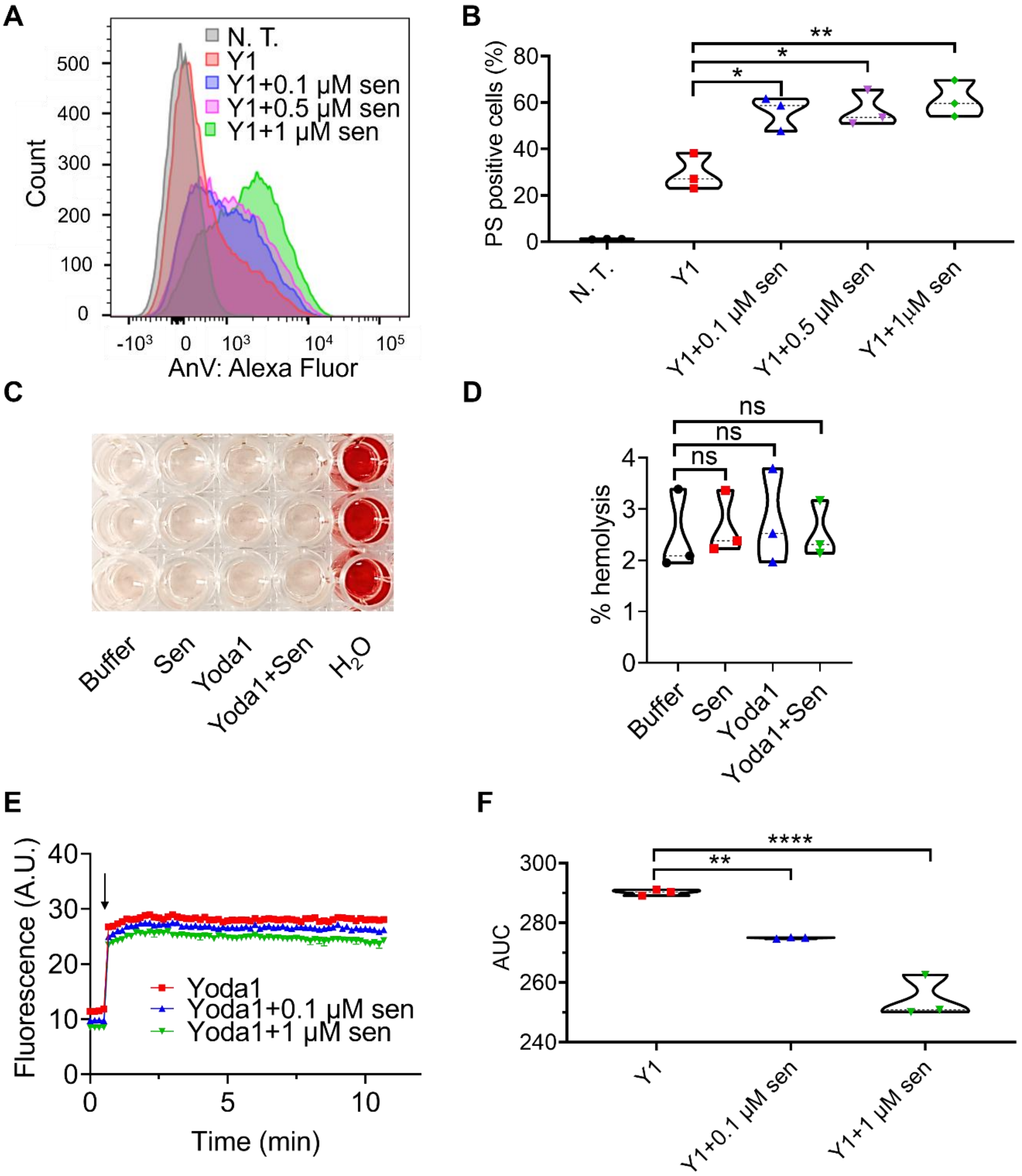


**Figure S8. Senicapoc paradoxically reduces $Ca^{2+}$ influx, but increases PS exposure in HD RBCs.** (A) Representative flow cytometry traces of RBC PS exposure after 10 minutes of 1 μM Yoda1 with different concentrations of Senicapoc, as measured by AnV binding. (B) Statistical analysis of 1 μM Yoda1-stimulated RBC PS exposure with dilutions of Senicapoc; one-way ANOVA, * $P$ = 0.0123, * $P$ = 0.0107, ** $P$ = 0.004 (n = 3). (C) Image of HD RBC hemolysis under different conditions in triplicate (rows): buffer, 1 μM Senicapoc, 2 μM Yoda1, and 1 μM Senicapoc +2 μM Yoda1. $H_2O$ was added to fully lyse the RBCs as a positive control (right). (D) Statistical analysis of HD RBC hemolysis in C. One-way ANOVA, ns (not significant) $P = 0.9799$, 0.9325, 0.999 for Sen, Yoda1, and Yoda1+Sen, respectively (n = 3). All hemolysis values were normalized by the $H_2O$ control. (E) $Ca^{2+}$ influx in HD RBCs measured by Calbryte 520 dye using a microplate reader. The arrow indicates stimulation of 1 μM Yoda1. (F) The AUC analysis of $Ca^{2+}$ influx in HD RBCs from panel E; one-way ANOVA ** $P = 0.0066$, **** $P < 0.0001$ (n = 3). N.T.: no treatment; sen: Senicapoc; Y1: Yoda1.
